# Beyond Generic Signal Peptides: ApexSP Enables Cargo-Specific Secretion Design

**DOI:** 10.64898/2026.07.29.741461

**Authors:** Chunyi Yang, Ruiyang Hou, Zhenyu Ma, Qiuyan Kang, ShuoJing Zuo, Limei Xu, Min Xiao, Xiuyun Wu, Xukai Jiang

## Abstract

Recombinant proteins are widely used in biopharmaceuticals, industrial manufacturing, and molecular diagnostics; however, efficient secretion remains a major bottleneck limiting their large scale production. As core elements controlling protein entry into secretion pathways, signal peptides do not function solely based on their own sequences, but rather depend on coordinated compatibility among cargo protein properties, secretion pathways, and host backgrounds. Current signal peptide engineering mainly relies on a limited number of commonly used signal peptides, empirical selection, and individual experimental screening, making it difficult to design efficient secretion elements tailored to specific expression systems. Here, we present ApexSP, a signal peptide design framework for secretion engineering tailored to individual cargo proteins. ApexSP is built upon a large scale, high quality signal peptide knowledge base and integrates a discrete diffusion generative model constrained by evolutionary information, a multitask biological filter incorporating topology information, and a signal peptide and mature protein compatibility ranking model to enable integrated design of signal peptides for specific expression systems. ApexSP achieved high accuracy in multiple attribute prediction tasks, including pathway classification and cleavage site prediction, reaching or exceeding the performance of existing signal peptide prediction tools. Moreover, the cargo protein aware ranking module improved the enrichment of candidates with high secretion performance. Experimental validation across multiple eukaryotic and prokaryotic expression systems demonstrated that screening only 10 candidate sequences generated by ApexSP yielded multiple designed signal peptides outperforming reference signal peptides, with the best performing design in the ApGA expressed Pichia pastoris system achieving a 1.71 fold increase in secretion performance compared with the α factor signal peptide. Overall, ApexSP advances signal peptide research from sequence prediction toward secretion element design, demonstrating that a single round of designed signal peptides can achieve high secretion performance suitable for further engineering optimization.

## 1. Introduction

Efficient secretion expression is essential for the large-scale production of recombinant proteins, including industrial enzymes, antibody fragments, cytokines, and vaccine antigens^1^. Compared with intracellular expression, secretion expression can reduce downstream purification complexity, decrease the risk of inclusion body formation, and facilitate proper folding and maturation of certain proteins^2,3^. As core secretion elements located at the N-terminus of secreted proteins, signal peptides direct precursor proteins into secretion pathways such as Sec and Tat, and influence transmembrane transport, signal peptide cleavage, mature protein N-terminal formation, and final extracellular accumulation^4^. Therefore, the selection of efficient signal peptides is a key challenge for improving heterologous protein secretion efficiency^5^.

Currently, signal peptide engineering mainly relies on a limited number of canonical signal peptides, empirical selection, and low-throughput experimental screening^6,7^. However, increasing evidence indicates that signal peptide function is not determined solely by its own sequence^8^. The same signal peptide may exhibit different secretion efficiencies when fused to different cargo proteins, while the same cargo protein with an identical signal peptide can also show different secretion outcomes under different host backgrounds, secretion pathways, or cleavage boundaries^9–11^. These observations indicate that there is no universally optimal signal peptide, and that so-called “strong signal peptides” often function only in specific cargo protein and host contexts. Therefore, signal peptide optimization should not be regarded simply as a screening problem for universal sequence elements, but rather as a combinatorial design problem jointly determined by signal peptides, mature proteins, secretion pathways, and host environments.

To address secretion engineering challenges, computational methods have been introduced to reduce experimental screening. SignalP^12^ and USPNet^13^ have significantly advanced signal peptide identification, Sec/Tat pathway classification, and cleavage-site prediction, enabling more accurate identification of signal peptides^14^. Meanwhile, several studies have begun to explore machine learning or generative models for designing new sequences with natural signal peptide characteristics^15,16^. However, generating sequences with native-like signal peptide features alone remains insufficient for secretion engineering applications. For practical applications, designed signal peptides only possess clear engineering value when they can efficiently direct secretion of specific cargo proteins and outperform commonly used universal signal peptides or conventionally screened candidates in experimental systems. Therefore, signal peptide design models should not only generate sequences satisfying Sec/Tat topology, cleavage-site constraints, and physicochemical characteristics, but also evaluate their compatibility with target mature proteins and enrich high-performing candidates in specific host systems^17^. However, due to the long-standing scarcity of large-scale signal peptide–cargo protein paired data and quantitative secretion labels^18^, a systematic design framework that simultaneously integrates signal peptide generation, biological constraint filtering, and cargo compatibility prediction remains lacking.

Here, we present ApexSP, a signal peptide design framework for cargo protein-tailored secretion engineering. We integrated and constructed a dataset containing millions of high-confidence signal peptide sequences. Based on this resource, ApexSP enables a systematic design process from signal peptide generation and functional constraint screening to cargo compatibility ranking. Experimental validation demonstrated that across multiple eukaryotic and prokaryotic expression systems, screening only 10 ApexSP-designed candidate sequences yielded multiple high-secretion candidates outperforming reference signal peptides. Overall, ApexSP advances signal peptide research from sequence identification and native-like sequence generation toward cargo protein-aware secretion element design, providing an effective strategy for improving heterologous protein secretion.

## 2. Materials and Methods

### 2.1 Dataset Construction and Preprocessing

To construct the signal peptide data resources used for ApexSP model training, validation, and performance evaluation, we integrated protein sequences from UniRef50, UniProtKB/Swiss-Prot, and several publicly available signal peptide datasets, including Signal6, USP_val, SPSDB, and SPE^16,18–21^. UniRef50 was primarily used to construct large-scale training sets of Sec- and Tat-type signal peptides, whereas UniProtKB/Swiss-Prot was used to supplement signal peptide–mature protein pairs with relatively high-quality annotations. Signal6_data, USP_val, and SPSDB were mainly used to validate the attribute prediction model and compare its performance with existing methods, while the SPE dataset was used for training or evaluating the signal peptide–mature protein compatibility model.

For the raw protein sequences, entries annotated as fragments were first removed to reduce the influence of incomplete sequences on signal peptide identification and cleavage site annotation. Subsequently, only protein sequences with lengths of 50–1000 aa and starting with methionine were retained for subsequent signal peptide mining and model training. To reduce data redundancy caused by highly similar sequences, the cleaned sequences were further clustered using CD-HIT at 95% sequence identity^22^, and representative sequences were retained as input for subsequent analyses.

### 2.2 Signal peptide annotation

To obtain high-confidence Sec and Tat signal peptide samples, we used USPNet and SignalP6 for two-stage annotation. First, USPNet was used to perform preliminary screening on the non-redundant protein sequences, and the sequences were classified into Sec signal peptide, Tat signal peptide, and non-signal peptide candidates. Subsequently, the Sec and Tat candidate sequences predicted by USPNet were separately input into SignalP6 for secondary validation, and the final predicted type as well as the N-region, H-region, and C-region topological regions were extracted from the SignalP6 output results. Only Sec or Tat sequences with consistent prediction results between USPNet and SignalP6 were retained as high-confidence signal peptide samples^12,13^. For each retained sequence, the protein ID, full-length protein sequence, pathway category, and corresponding N-region, H-region, and C-region boundaries were recorded for subsequent generative models, biological filters, and regional physicochemical feature analyses.

To construct a non-signal peptide control set, we collected protein sequences annotated as intracellularly localized from UniProtKB/Swiss-Prot and preprocessed them using the same sequence cleaning and redundancy removal procedure. Candidate non-signal peptide sequences were first screened as the NO_SP category by USPNet and then further validated as the NO_SP category by SignalP6. Only sequences simultaneously recognized as non-signal peptides by USPNet and SignalP6 were retained as high-confidence NO_SP samples. Since this category does not contain definable signal peptide topological regions, its N-region, H-region, and C-region annotations were all marked as null values.

In addition to signal peptide pathway and topology annotations, sequences derived from UniRef50 were further annotated according to their taxonomic origin. Based on the TaxID information provided in the FASTA headers^23^, each sequence was mapped to the corresponding node in the NCBI Taxonomy database, and its taxonomic lineage was traced upward to the superkingdom level. Sequences assigned to Eukaryota were labeled as eukaryotic, whereas those assigned to Bacteria or Archaea were labeled as prokaryotic. Sequences with unresolvable TaxIDs or belonging to other categories were excluded from the taxonomic-origin supervised training set. Following these procedures, the ApexSP dataset contained signal peptide pathway labels, taxonomic-origin labels, and topology labels predicted by SignalP 6, thereby supporting Sec/Tat/ NO_SP classification, eukaryotic/prokaryotic origin identification, N/H/C-region learning, and signal peptide generative model training.

### 2.3 Evolutionarily constrained discrete diffusion model for signal peptide generation

Based on the Sec and Tat signal peptide samples described above, ApexSP constructed a pathway-conditioned discrete denoising diffusion probabilistic model (D3PM)^24,25^ to learn the sequence distributions of signal peptides from different secretion pathways. The model vocabulary consisted of 20 standard amino acid tokens and one PAD token. For each training sequence, the signal peptide fragment was extracted according to the C-region end predicted by SignalP 6 and encoded to a maximum length of 70 amino acids. Shorter sequences were padded at the C terminus with PAD tokens, and the padded positions were not treated as valid amino acids during the interpretation of the generated sequences.

The diffusion process was defined in the discrete amino acid space. Given a native signal peptide sequence x0, the model gradually perturbed amino acid identities through a forward transition matrix at a random time step t to generate a noisy sequence xt. The noise intensity used a cosine schedule^26^, and the number of diffusion steps was set to 100. Different from uniformly random replacement noise, ApexSP introduced the BLOSUM62 amino acid substitution matrix as an evolutionary prior in the model. The BLOSUM62 matrix^27^ was converted into an amino acid substitution probability matrix after temperature scaling, making the forward diffusion more likely to perturb amino acids among evolutionarily substitutable residues. This design introduces the evolutionary substitution rules of protein sequences into the discrete diffusion process to reduce the destruction of amino acid semantic structure caused by completely random noise.

The denoising network takes the noisy sequence xt, diffusion time step t, and pathway label as inputs to predict the original amino acid identities. Amino acid tokens were first embedded into continuous vectors, followed by a one-dimensional convolution module to extract local sequence features. Positional embeddings, timestep embeddings, and pathway condition embeddings were then added. The diffusion timestep t was encoded using sine and cosine functions and projected into a timestep representation, whereas pathway labels were represented by learnable embeddings to enable pathway-conditioned generation. The fused sequence representations were subsequently processed by a four-layer Transformer encoder to model long-range sequence dependencies^28^, followed by a convolutional decoder that output the amino acid probability distribution at each sequence position.

The model was trained using all processed signal peptide samples. The training objective was to reconstruct the native sequence x0x_0x0 from the noisy sequence xtx_txt, and amino acid reconstruction cross-entropy was used as the optimization objective. Training was performed using the AdamW optimizer with a learning rate of 1×10−41 \times 10^{-4}1×10−4, a batch size of 4096, and 10 training epochs. To evaluate the contribution of different design modules, systematic ablation experiments were performed. The baseline model did not include positional embeddings, BLOSUM62 evolutionary noise, C-terminal site weighting, pathway condition embeddings, or Tat RR motif weighting. Based on the baseline, five modules were evaluated sequentially: (A) positional embeddings; (B) BLOSUM62 evolutionary noise; (C) C-terminal −1/−3 site weighting; (D) Sec/Tat pathway condition embeddings; and (E) Tat RR motif weighting. A total of nine models were trained, including the baseline, A, A+B, A+C, A+D, A+B+D, A+B+C, A+B+C+D, and A+B+C+D+E. All ablation models were trained using the same data source and major hyperparameters, and were compared using amino acid Top-5 reconstruction accuracy, Tat RR motif recovery, and recovery accuracy at the −1 and −3 cleavage-site positions^29^.

### 2.4 Candidate signal peptide generation

After training the discrete diffusion model, ApexSP generated candidate signal peptide sequences through reverse diffusion sampling. Sampling was initialized from random discrete amino acid noise and proceeded through iterative denoising conditioned on a specified Sec or Tat pathway label. At each reverse diffusion step, the model output an amino acid probability distribution at each position based on the current sequence state and timestep, from which the next sequence state was sampled^30^. During decoding, sequence generation was terminated upon encountering a PAD token.

To reduce low-complexity fragments and homopolymeric amino acid repeats^31,32^, a consecutive amino acid repetition-blocking strategy was applied during sampling. When three consecutive identical amino acids had already occurred immediately before a given position, the sampling probability of that amino acid at the current position was reduced to 10^-10^. The probabilities of the remaining amino acids were then renormalized before sampling continued^33^.

### 2.5 Signal peptide feature characterization

To determine whether the generated sequences retained the regional organization of natural signal peptides, natural Sec and Tat signal peptides were used as references. The boundaries of the N-region, H-region, and C-region of natural signal peptides were determined by the SignalP6 topological region annotation. The total length of a signal peptide was defined as the C-region end position, namely the length of the N-terminal fragment before the predicted cleavage site. For natural Sec and Tat signal peptides, we separately calculated the lengths of the N-region, H-region, and C-region, and divided them into three categories according to pathway and species origin: prokaryotic Tat, prokaryotic Sec, and eukaryotic Sec.

To compare regional physicochemical patterns among signal peptides of different lengths, the N-region, H-region, and C-region of each signal peptide were mapped to 4, 8, and 4 fixed dimensions, respectively. Each region was divided into a fixed number of intervals according to its length, and the average amino acid charge, hydrophobicity, and residue volume were calculated within each interval. This fixed-dimensional regional representation was used to compare the relative physicochemical patterns among different sequence sets, but did not represent the absolute distribution of residue-by-residue positions in the original sequence. The physicochemical properties of amino acids were calculated using fixed lookup tables, including Kyte-Doolittle hydrophobicity, amino acid volume, and charge state^34,35^. N-region charge, H-region hydrophobicity, and C-region residue volume were used as representative features of the regional organization of signal peptides to evaluate whether the generated sequences retained the classical signal peptide features of a positively charged N-region, hydrophobic H-region, and small-residue C-region.

### 2.6 Quality evaluation of generated signal peptides

To systematically evaluate the quality of signal peptides generated by the discrete diffusion model, 10,000 candidate sequences were sampled under each of the Sec and Tat conditions, yielding a total of 20,000 generated signal peptides. The generated sequences were evaluated in terms of length distribution, regional physicochemical properties, amino acid composition, C-terminal residue preference, the Tat twin-arginine motif, nearest-neighbor similarity, internal diversity, and model perplexity^36^.

The length distributions of the generated sequences were compared with those of the natural reference sequences separately for the Sec and Tat classes. The length of a natural signal peptide was defined by the C-region end position predicted by SignalP 6, corresponding to the N-terminal segment preceding the predicted cleavage site. The length of a generated sequence was defined as the number of non-PAD amino acids obtained after reverse diffusion sampling and decoding. Length distributions were visualized using grouped raincloud plots. To avoid excessive point density, no more than 800 sequences were randomly sampled from each group for scatter visualization, whereas the complete sequence sets were used for distributional analyses.

Regional physicochemical properties of the generated and natural reference sequences were compared using the procedure described in Section 2.4. For both natural and generated sequences, N-, H-, and C-region boundaries were predicted using SignalP 6, and the distributions of N-region charge, H-region hydrophobicity, and C-region residue volume were compared.

Cleavage-site preferences of the generated sequences were evaluated based on their C-terminal tripeptide composition. For each generated signal peptide, the final three amino acids were extracted, and the occurrence probabilities of high-frequency tripeptide motifs were calculated separately for the Sec and Tat sequence sets. This analysis was used to determine whether the generated sequences were enriched in combinations containing small residues such as Ala and Ser, which are characteristic of the C-terminal region of cleavable signal peptides.

To determine whether the generated Sec and Tat sequences occupied distinguishable pathway-associated sequence spaces, their latent sequence structure was analyzed using k-mer features. Each signal peptide was represented as an overlapping 3-mer count vector and subjected to L2 normalization. The resulting vectors were reduced in dimensionality using TruncatedSVD and subsequently visualized with UMAP.

The fidelity of the generated sequences was evaluated using nearest-neighbor similarity^37^. For each generated sequence, the most similar sequence in the natural reference set was identified, and sequence identity was used as the nearest-neighbor similarity score. To improve the efficiency of large-scale comparisons, a 5-mer-based inverted index was first constructed, and nearest-neighbor searches were restricted to candidate sequences differing in length by no more than 8 amino acids. For each generated sequence, up to 1,000 candidate natural sequences were retained for exact similarity calculation.

Generated-sequence diversity was evaluated using internal sequence distances within length-stratified groups^38^. Sequences were divided into multiple length intervals, and sequence pairs were randomly sampled within each interval. Pairwise distance was defined as 1−sequence identity. Up to 1,000 sequence pairs were sampled from each length interval to estimate mean internal diversity. This metric was used to assess whether the model exhibited mode collapse and whether it generated diverse candidate signal peptides across different length ranges.

Functional motifs in Tat-conditioned sequences were evaluated by analyzing the RR motif. For each Tat sequence, the position of the first consecutive twin-arginine motif was identified^39^, and both its absolute position and its position relative to sequence length were recorded. Sequences were then grouped by length, and the mean position of the first RR motif and its standard error were calculated for each group. This analysis assessed whether the generated Tat signal peptides retained the RR motif near the N terminus rather than displaying an abnormal downstream shift with increasing sequence length.

Model perplexity was used to evaluate prediction confidence for signal peptides of different lengths^40^. Sec and Tat test sequences were grouped into six length intervals: 0–15, 15–20, 20–25, 25–30, 30–35, and 35–70 amino acids. For each group, amino acid prediction cross-entropy was calculated at multiple diffusion noise timesteps and converted to perplexity. Lower perplexity indicated greater confidence in amino acid recovery. This analysis was used to evaluate the stability of the model across signal peptides of different lengths and secretion pathways.

### 2.7 Topology-aware multi-task biological filter

To biologically screen the generated candidate sequences, ApexSP constructed a topology-aware multitask model that simultaneously predicted signal peptide pathway, residue-level topology, and taxonomic origin. The model comprised three tasks: T1, sequence-level pathway classification into NO_SP, Sec, and Tat; T2, residue-level topology annotation into the N-region, H-region, C-region, N-terminal region of the mature protein, and PAD; and T3, sequence-level taxonomic-origin classification into eukaryotic and prokaryotic classes. All input sequences were truncated or padded to a length of 70 amino acids. To accommodate different input formats, the training samples consisted of complete signal peptide sequences, 70-aa N-terminal fragments of secreted proteins, and randomly truncated N-terminal fragments at proportions of 60%, 30%, and 10%, respectively. The model used ESM2_t33_650M_UR50D^41^ as the backbone encoder and was parameter-efficiently fine-tuned using DoRA adapters. DoRA modules were introduced into the query, key, value, attention output dense, and intermediate dense projections of each Transformer layer, with the rank, alpha, and dropout set to 16, 32, and 0.1, respectively^42^. For each residue, the ESM2 representation was concatenated with hydrophobicity, residue volume, charge, and normalized positional features. The concatenated features were sequentially processed by linear projection, layer normalization, GELU activation, and dropout to obtain a shared residue representation. T1 and T3 used masked mean pooling to derive sequence-level representations, which were then passed through multilayer perceptrons to output pathway and taxonomic-origin probabilities. For T2, the shared representation was first projected to 640 dimensions, followed by one Transformer encoder layer and two one-dimensional convolutional layers to capture contextual information and local boundary features. A linear classification layer and a conditional random field were then used to output residue-level topology labels^43^.

The model was trained using joint multitask optimization^44^. Weighted cross-entropy losses were used for T1 and T3, whereas T2 was optimized using the conditional random field negative log-likelihood. The total loss was defined as the sum of the three task-specific losses. Higher weights were assigned to the N-, H-, and C-region labels in the T2 loss, with the highest weight assigned to the C-region, whereas lower weights were assigned to the mature-protein N-terminal and PAD labels. Training was performed using the AdamW optimizer with both the learning rate and weight decay set to 1×10^−4^, a batch size of 8, and gradient accumulation. The model was trained for up to four epochs with an early-stopping patience of three, and the checkpoint with the lowest validation loss was retained. To assess the contributions of ESM2 pretrained representations and residue-level topology supervision, four ablation models were evaluated: a model without ESM2 or T2, a model with T2 but without ESM2, a model with ESM2 but without T2, and the complete model containing both ESM2 and T2. All models used the same data splits, training strategy, and evaluation metrics.

During inference, the model simultaneously output pathway probabilities, taxonomic-origin probabilities, residue-level topology labels, and cleavage-site positions. The T1 and T3 classes were determined by the highest softmax probability, whereas the T2 labels were obtained by conditional random field decoding. For sequences predicted as Sec or Tat, the cleavage site was defined as the position immediately following the final residue of the predicted C-region. For sequences predicted as NO_SP, no N-, H-, or C-region or cleavage site was assigned. Generated sequences were subsequently filtered according to agreement with the target pathway, topology completeness, and predicted taxonomic origin.

Model performance was evaluated on the internal test set, an independent UniRef50 validation set, and publicly available benchmark datasets. For the internal test set, accuracy, precision, recall, F1 score, and ROC–AUC were calculated for pathway classification, topology annotation, and taxonomic-origin classification. For pathway classification, one-versus-rest ROC curves were generated for the NO_SP, Sec, and Tat classes. UniRef50 validation sequences were divided into five groups according to signal peptide length: 15–24, 25–34, 35–44, 45–54, and ≥55 amino acids, and pathway- and taxonomic-origin classification performance was evaluated separately for each group. Topology prediction was further assessed using the localization errors of the N/H, H/C, and C-region terminal boundaries. For comparison with existing methods, SignalP 6 and USPNet were applied to the independent validation set, and all predictions were mapped to the common classes NO_SP, Sec, and Tat. Performance was compared separately across archaeal, eukaryotic, Gram-negative bacterial, and Gram-positive bacterial subsets. Predicted cleavage sites from ApexSP, SignalP 6, and USPNet were converted to residue positions relative to the N terminus of the input sequence, and localization accuracy was calculated at error tolerances of 0, ≤1, ≤3, and ≤5 residues.

### 2.8 Signal peptide–mature protein compatibility ranking model

To prioritize candidate signal peptides for a specific cargo protein^45,46^, a signal peptide–mature protein compatibility ranking model was constructed. Candidate signal peptide sequences and the N-terminal sequences of the downstream mature proteins were used as two separate input branches, and a continuous compatibility score was generated for relative ranking of different candidate signal peptides within the same mature protein context.

To evaluate the effect of mature protein input length on ranking performance, six N-terminal truncation lengths of 3, 10, 50, 100, 200, and 500 aa were tested. The maximum signal peptide input length was fixed at 70 aa. All sequences were truncated or padded at the C terminus to the predefined input length, and padding positions were excluded from feature aggregation using attention masks.

A dual-branch encoding architecture with latent-space matching was used. Amino acid sequences were first converted into discrete tokens and mapped to 128-dimensional trainable embeddings. Each residue embedding was further combined with normalized Kyte–Doolittle hydrophobicity, residue volume, and charge features. Independent positional embeddings were used for the signal peptide and mature protein branches. An 8-dimensional Sec/Tat pathway embedding was concatenated with the sequence-level representation from each branch.

The two branches used structurally identical but parameter-independent encoders. Each encoder contained two Transformer encoder layers followed by two one-dimensional convolutional modules with kernel sizes of 5 and 3, respectively. Each convolutional module consisted of one-dimensional convolution, batch normalization, GELU activation, and dropout. Cross-branch feature interaction was subsequently implemented using multi-head cross-attention, with the mature protein representation used as the query and the signal peptide representation used as the key and value. The attention output was integrated into the mature protein branch through a residual connection.

The residue-level representations after cross-branch interaction were aggregated by masked mean pooling. The signal peptide and mature protein representations were separately concatenated with the pathway embedding and projected into 32-dimensional latent representations through independent projection layers. During pretraining, the cosine similarity between the two projected representations was used as the predicted compatibility score. During fine-tuning, the two projected representations were concatenated and passed to a multilayer perceptron prediction head to produce a continuous compatibility score.

### 2.9 Pseudo-label pretraining and fine-tuning using experimental secretion data

Because large-scale quantitative secretion data for signal peptide–mature protein combinations remain limited, pseudo-label pretraining was first performed to initialize the compatibility model. The pretraining data were derived from the high-confidence natural secreted protein dataset constructed for ApexSP ^19^. Based on the predicted end position of the signal peptide C-region, each full-length protein sequence was separated into an N-terminal signal peptide and a downstream mature protein. The N-terminal fragment of the mature protein was then truncated to the predefined input length.

Each natural signal peptide–mature protein pair was expanded into four pseudo-labeled sample types^47^: the native signal peptide paired with its original mature protein; a signal peptide containing 10% conservative amino acid substitutions paired with the original mature protein; a signal peptide containing 30% random amino acid substitutions paired with the original mature protein; and a negative sample in which the signal peptide was replaced by a length-matched placeholder sequence. Target compatibility labels for these four sample types were sampled from normal distributions with means of 0.93, 0.78, 0.40, and 0.05, respectively. Conservative substitutions were introduced according to physicochemical similarity groups, including small or polar residues, basic residues, acidic or amide-containing residues, aromatic residues, and hydrophobic aliphatic residues.

During pretraining, the signal peptide sequence, mature protein N-terminal sequence, and Sec/Tat pathway label were jointly provided as model inputs. The cosine similarity between the projected representations of the two branches was regressed against the pseudo-label compatibility score. The model was optimized using mean squared error loss and the AdamW optimizer. Data were split according to the original protein identifier to ensure that pseudo-labeled samples derived from the same parent protein were assigned to the same data subset. Early stopping was applied, and the model parameters corresponding to the lowest validation loss were retained.

Following pseudo-label pretraining, supervised fine-tuning was performed using the AmyQ– *Bacillus subtilis* secretion system in the SPE dataset^16^. The SP, MP, and WA fields were used as the signal peptide sequence, mature protein sequence, and experimentally measured secretion yield, respectively, and the normalized WA_true value was used as the regression target. Samples lacking SP, MP, or yield annotations, as well as sequences containing non-standard amino acids, were removed. All samples were encoded as Sec-pathway sequences.

For fine-tuning, the pretrained model parameters were used for initialization, and the secretion-yield prediction head was trained using the 32-dimensional projected representations from the two branches. The dataset was divided into training, validation, and test sets using a fixed random seed, with proportions of 70%, 20%, and 10%, respectively. A model with the same architecture and data split but random parameter initialization was trained in parallel as a pretraining ablation control. The original SPE dataset includes corrections to the SP and MP sequences based on the actual cleavage boundary of each signal peptide^16^. During fine-tuning, the corrected cleavage boundaries were directly used to reconstruct the SP–MP input pairs, thereby increasing the sequence diversity of the mature protein inputs.

### 2.10 Compatibility model evaluation and external validation

To assess the contributions of individual components of the compatibility model, systematic ablation experiments were performed by comparing model configurations that differed in mature protein N-terminal input length, ESM2 pretrained representations, the cross-attention module, and the pseudo-label pretraining strategy. All other network architectures, data splits, and training parameters were kept unchanged across ablation settings.

Model performance was evaluated in terms of correlation, within-group ranking quality, and enrichment of high-secretion candidates. The evaluation metrics included Spearman’s correlation coefficient, pairwise accuracy, Top-1 hit rate, NDCG@10, NDCG@20, and EF@10%^48^. In addition, random-ranking baselines were constructed by shuffling the predicted scores 100 times and were evaluated using the same metrics.

External validation was performed using multiple independent, publicly available secretion datasets covering hosts including *Bacillus subtilis*, *Corynebacterium glutamicum*, and *Lactiplantibacillus plantarum*, as well as cargo proteins including cutinase, nuclease, and alkaline xylanase^18^. Signal peptide and mature protein sequences from each dataset were processed using the same procedure as that applied during model training and were then input into the compatibility model to obtain prediction scores. Because the datasets differed in host background, experimental assay, and secretion-yield range, prediction scores were used only for relative ranking within each dataset. Performance was evaluated using within-dataset ranking correlations, percentile ranks, and enrichment metrics for high-secretion candidates.

### 2.11 Experimental validation of the eukaryotic system

Glucoamylase ApGA from *Aureobasidium pullulans* was selected as the eukaryotic cargo protein, and *Pichia pastoris* X-33 was used as the heterologous expression host^49^. The strains used in this study are listed in Table S1. The first 35 amino acid residues of ApGA were identified as its native signal peptide. After removal of this region, the mature protein sequence was inserted downstream of the Kex2 cleavage site of the α-factor leader in the pPICZαA vector. The recombinant expression construct was synthesized by Sangon Biotech (Shanghai, China), and the codon-optimized DNA sequence used for gene synthesis is provided in Table S2. The recombinant plasmid was first transformed into *Escherichia coli* DH5α, and transformants were selected on LB agar containing 50 μg/mL Zeocin. Positive clones were verified by sequencing to obtain the recombinant ApGA expression plasmid.

The verified recombinant plasmid was introduced into *P. pastoris* X-33 by electroporation, and transformants were selected on YPD agar containing 100 μg/mL Zeocin. Sequence-verified transformants were inoculated into 5 mL YPD medium containing 100 μg/mL Zeocin and cultured at 30°C and 220 rpm for 24 h. The resulting culture was transferred into 50 mL YPD medium at an inoculation ratio of 1% (v/v) and cultured at 30°C and 220 rpm until the OD_600_ reached 2–4. Methanol was then added to a final concentration of 1% for induction. Induction was continued for 72 h, with methanol replenished every 24 h. After fermentation, the culture was centrifuged at 8,000 rpm for 10 min at 4°C, and the supernatant was collected as the crude enzyme preparation.

Secreted ApGA was detected by SDS–PAGE, and glucoamylase activity in the fermentation supernatant was measured using the 3,5-dinitrosalicylic acid method. One unit of enzyme activity was defined as the amount of enzyme required to produce 1 μg of reducing sugar per minute per milliliter of fermentation supernatant.

For the ApGA mature protein, 100,000 candidate signal peptides were generated under the Sec condition. The generated sequences were sequentially processed using the biological filter and the signal peptide–mature protein compatibility ranking model, and 10 candidates were selected from the top-ranked sequences for experimental validation (Table S3). These candidate sequences were used to replace the signal peptide region of the α-factor leader, and the primers used are listed in Table S4. The resulting recombinant strains were fermented under the conditions described above, and ApGA secretion was evaluated by SDS–PAGE and enzyme activity assays^50^.

### 2.12 Experimental validation of the prokaryotic system

Protease AprE2 from *Bacillus clausii* was selected as the prokaryotic cargo protein, and *Bacillus subtilis* WB600 was used as the expression host. Based on sequence alignment against the NCBI database, the signal peptide from the protein showing the highest sequence similarity to AprE2 (UniProt: Q99405) was used as the reference Sec signal peptide, whereas the PhoD signal peptide was used as the reference Tat signal peptide. The target protein sequence was fused separately to the two reference signal peptides and cloned into the pP43NMK vector to generate the pP43NMK-Sec-aprE2 and pP43NMK-Tat-aprE2 expression plasmids.

The recombinant plasmids were first transformed into *E. coli* DH5α, and transformants were selected on LB agar containing 100 μg/mL ampicillin after incubation at 37°C for 12–16 h. After sequence verification, the recombinant plasmids were transformed into *B. subtilis* WB600^51^. Expression strains were selected on LB agar containing 50 μg/mL kanamycin after incubation at 37°C for 12 h. Sequence-verified strains were inoculated into 1 mL LB medium containing 50 μg/mL kanamycin and cultured at 37°C and 220 rpm for 12 h. The resulting seed cultures were transferred into 50 mL LB medium at an inoculation ratio of 5% (v/v) and cultured at 37°C and 220 rpm for 24 h. The fermentation cultures were centrifuged at 10,000 rpm for 10 min at 4°C, and the supernatants were collected for subsequent analyses.

Secreted AprE2 was detected by SDS–PAGE. For the protease activity assay, 100 μL of 2% (w/v) casein substrate was mixed with 100 μL of fermentation supernatant and incubated at 40°C for 10 min. After centrifugation at 10,000 rpm for 10 min, 100 μL of the resulting supernatant was mixed with 500 μL of 0.4 M sodium carbonate and 100 μL of Folin–Ciocalteu reagent. The reaction was incubated at 40°C for 20 min, and absorbance was measured at 663 nm.

For the AprE2 mature protein, 100,000 candidate signal peptides were generated under each of the Sec and Tat conditions. The generated sequences were sequentially processed using the biological filter and the signal peptide–mature protein compatibility ranking model, and 10 candidates were selected from the top-ranked sequences for each pathway for experimental validation. The candidate signal peptides were used to replace the corresponding reference Sec or Tat signal peptide^52^. Their amino acid sequences and the primers used are listed in Tables S3 and S4, respectively. The resulting recombinant strains were fermented under the conditions described above, and AprE2 secretion was evaluated by SDS–PAGE and protease activity assays.

## 3. Results

### 3.1 ApexSP enables cargo protein-aware signal peptide design through an integrated framework

To enable cargo protein–specific signal peptide design, we developed ApexSP, an end-to-end design framework integrating large-scale signal peptide knowledge mining, sequence generation, biological filtering, and cargo protein compatibility prediction (Fig. 1a).

**Figure 1.**
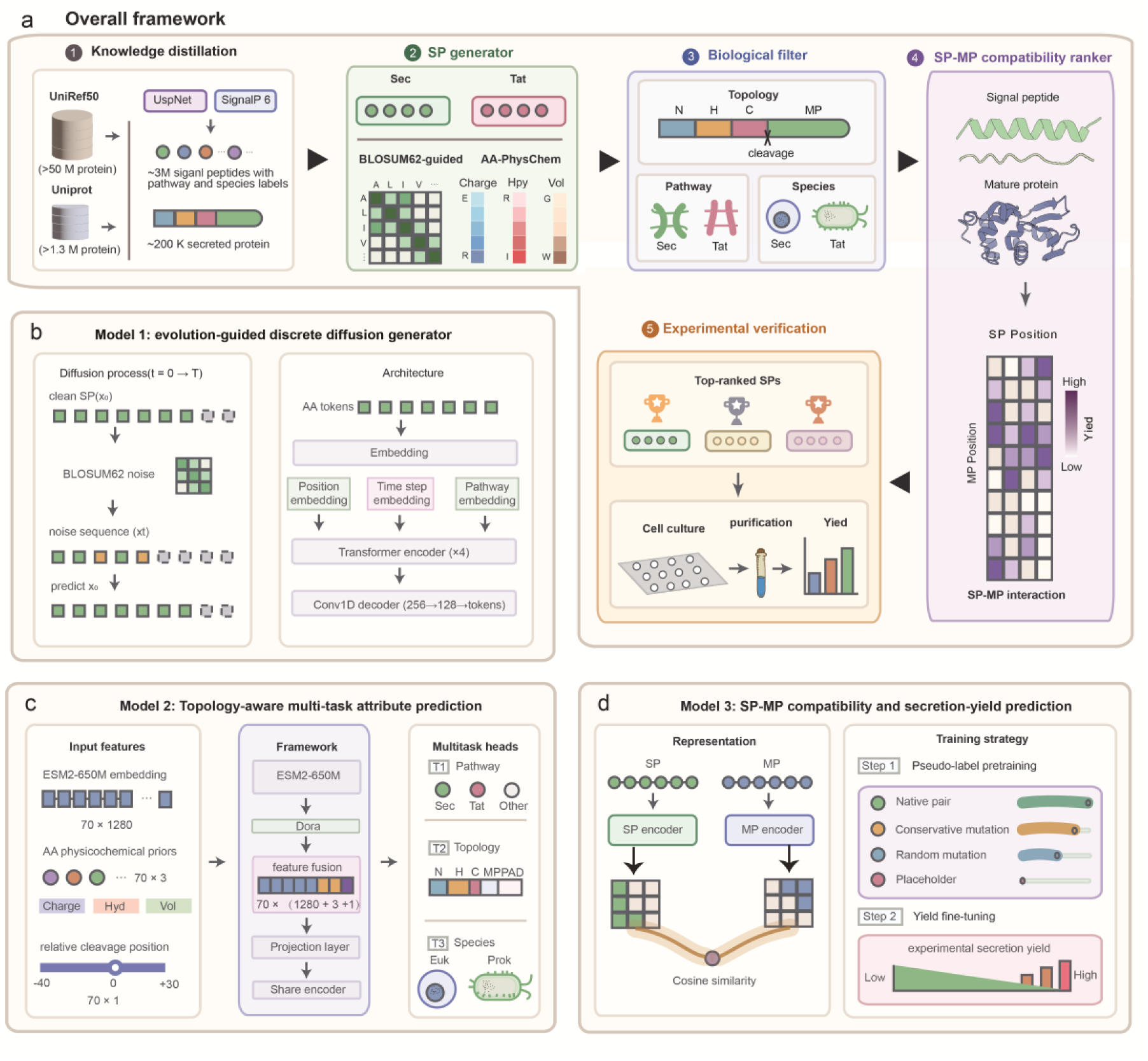
ApexSP framework and core model design. **(a)** Overview of the ApexSP framework integrating signal peptide knowledge distillation, sequence generation, biological filtering, SP–MP compatibility ranking, and experimental validation. **(b)** Evolution-guided discrete diffusion model for conditional generation of Sec and Tat signal peptides using BLOSUM62-informed amino acid substitution constraints. **(c)** Topology-aware multi-task model for biological filtering, jointly predicting secretion pathway, topology regions, and species origin. **(d)** SP–MP compatibility model based on dual-tower representation learning, enabling cargo protein-aware ranking of candidate signal peptides and prediction of secretion potential.

Large-scale protein sequence resources were integrated and consistently annotated using SignalP6 and USPNet to establish a high-confidence signal peptide knowledge base. Specifically, ApexSP development incorporated multiple data resources, including UniRef50, UniProtKB/Swiss-Prot, Signal6_Set, SPSDB, and SPE (Table S5). Among these datasets, UniRef50 provided the largest-scale signal peptide resource for model learning, containing 3.57 million Sec-type and 74 thousand Tat-type signal peptides after filtering. UniProtKB/Swiss-Prot further provided more than 200,000 high-quality signal peptide samples and signal peptide–mature protein paired information for cargo protein compatibility modeling. In addition to secretion pathway annotation, all source sequences were further linked to NCBI Taxonomy information and classified as prokaryotic or eukaryotic according to their taxonomic hierarchy. To assess data independence between model training and external validation, sequence overlap analyses were systematically performed between training datasets and external validation datasets (Table S6), revealing minimal overlap among most datasets.

Based on the aforementioned body of knowledge, ApexSP was constructed as a cascade design framework consisting of three interconnected modules. The generation module learns the sequence distribution of natural Sec/Tat signal peptides and produces candidate secretion signals (Fig.1b). The biological filtering module further constrains generated candidates based on secretion pathway, topology, and species origin (Fig.1c). The compatibility ranking module prioritizes candidate sequences by leveraging signal peptide–mature protein relationships (Fig.1d). Through sequential optimization across these modules, ApexSP enables the identification of cargo-adapted signal peptides from a large generated sequence space.

### 3.2 ApexSP-generated signal peptides exhibit a clear secretory signal peptide generation preference

To identify the key factors influencing signal peptide generation quality, different model designs were first compared through a denoising reconstruction task. Compared with the baseline model incorporating only positional embeddings, introducing BLOSUM62-based evolutionary noise improved amino acid recovery accuracy at the −1 and −3 positions upstream of the cleavage site from 0.6786 and 0.6812 to 0.7327 and 0.7363, respectively. The Top-5 amino acid reconstruction accuracy increased from 0.9522 to 0.9744, and the recovery rate of the Tat twin-arginine motif increased from 0.9028 to 0.9384 (Table S7). In contrast, introducing C-terminal position weighting or pathway-specific conditional embeddings alone resulted in only limited improvements. These results indicate that the evolutionary substitution prior is the primary factor contributing to improved recovery of both critical positions and overall sequence reconstruction capability.

Subsequently, high-confidence natural signal peptide datasets were statistically analyzed to establish reference characteristics for regional organization and pathway-specific features. Prokaryotic Tat signal peptides exhibited longer N- and H-regions, with average lengths of approximately 11 and 17 aa, respectively. In comparison, eukaryotic Sec signal peptides showed average N- and H-region lengths of approximately 4 and 13 aa, while prokaryotic Sec signal peptides exhibited lengths of approximately 5 and 14 aa, respectively. The C-regions of all three signal peptide classes showed similar lengths of approximately 5 aa, indicating that length constraints near the cleavage boundary are relatively conserved (Fig. 2a). After length normalization, all three signal peptide classes displayed typical regional physicochemical organization: the N-region showed a positive charge tendency, the H-region maintained high hydrophobicity, whereas the C-region near the cleavage site exhibited substantially reduced hydrophobicity and residue volume (Fig. 2b). Sequence logos further revealed enrichment of the RR motif in the N-region of Tat signal peptides, enrichment of Lys and Arg residues in prokaryotic Sec signal peptides, and a preference for small-volume residues such as Ala and Ser within the C-region of all signal peptide classes (Fig. 2c).

**Figure 2.**
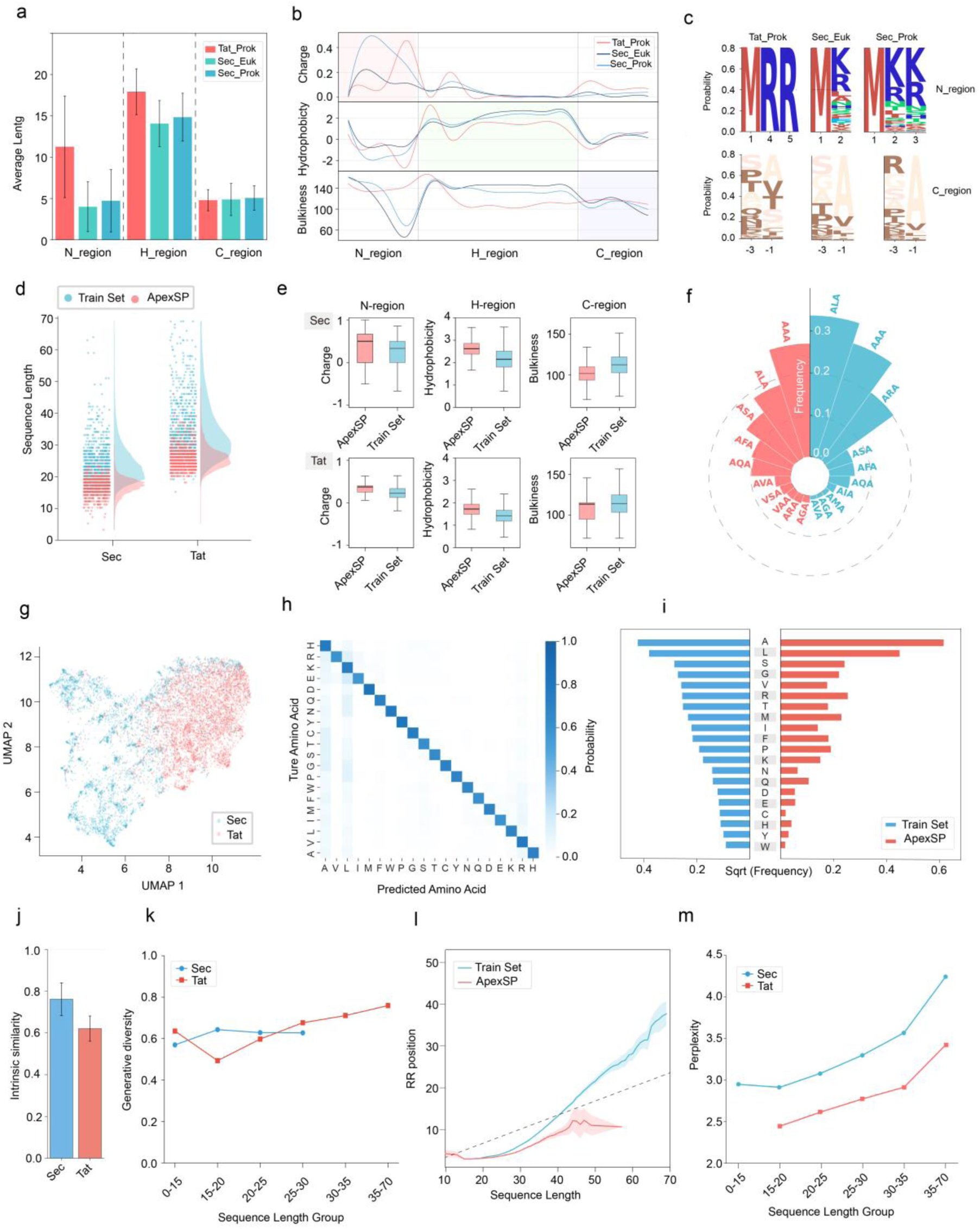
ApexSP generates signal peptides with canonical secretion features. **(a)** Length distribution of N-, H-, and C-regions in high-confidence natural signal peptides. **(b)** Region-specific physicochemical properties of natural signal peptides, including charge, hydrophobicity, and residue bulkiness. **(c)** Sequence logo analysis of N- and C-regions from Tat-prokaryotic, Sec-eukaryotic, and Sec-prokaryotic signal peptides. **(d)** Length distribution comparison between ApexSP-generated signal peptides and natural training sequences. **(e)** Comparison of region-specific physicochemical properties between ApexSP-generated and natural signal peptides. **(f)** Tripeptide composition frequencies around cleavage sites of ApexSP-generated signal peptides. **(g)** UMAP visualization of generated Sec and Tat signal peptides in latent representation space. **(h)** Amino acid prediction confusion matrix of the generation model. **(i)** Comparison of amino acid composition between ApexSP-generated sequences and natural training sequences. **(j)** Intrinsic similarity between ApexSP-generated sequences and natural signal peptides. **(k)** Sequence diversity of generated signal peptides across different length groups. **(l)** Positional distribution of the RR motif in Tat signal peptides across different sequence lengths. **(m)** Perplexity of ApexSP-generated sequences across different length groups.

Based on these reference features, ApexSP-generated sequences largely recapitulated the characteristics of natural signal peptides. Generated Sec sequences were mainly distributed within 12–25 aa, whereas Tat sequences were mainly distributed within 22–35 aa, preserving the overall tendency of Tat signal peptides to be longer than Sec signal peptides. Compared with the training dataset, generated sequences displayed a more concentrated length distribution, with a reduced long-sequence tail (Fig. 2d). In terms of regional physicochemical properties, both generated Sec and Tat signal peptides exhibited increased positive charge in the N-region and enhanced hydrophobicity in the H-region, together with reduced residue volume in the C-region, indicating stronger representation of the classical regional physicochemical features of signal peptides (Fig. 2e). In addition, the C-terminal regions of generated sequences were enriched in tripeptides composed of small-volume residues, with Ala-rich combinations such as ALA and AAA occurring at high frequencies (Fig. 2f), suggesting that residue preferences near the cleavage region were also preserved.

Sequence space analysis further showed that generated Sec and Tat sequences formed continuous yet distinguishable distributions in the UMAP space, indicating that the model preserved pathway-specific differences while maintaining shared global signal peptide characteristics (Fig. 2g). The amino acid recovery matrix exhibited clear diagonal enrichment, with major off-diagonal signals concentrated between residues with similar physicochemical properties, consistent with the conservative substitution patterns guided by BLOSUM62 (Fig. 2h). The overall amino acid composition of generated sequences followed the same trend as the natural datasets (Fig. 2i). Ala represented the most frequent residue (sqrt frequency 0.6 in generated sequences versus 0.45 in the training set), while Leu, Ser, Gly, Val, and Arg remained highly abundant, and low-frequency residues such as Trp, Tyr, and His remained rare.

Generated sequences also maintained a balance between sequence fidelity and diversity. The nearest-neighbor similarity of generated Sec and Tat sequences relative to natural reference datasets was 0.76 and 0.62, respectively (Fig. 2j), indicating that the generated sequences remained within the natural sequence space rather than simply reproducing training examples. Internal diversity across different length ranges remained between 0.49 and 0.76, with no obvious evidence of mode collapse (Fig. 2k).

Finally, to evaluate the preservation of local functional constraints and overall generation stability, the position of the Tat-specific RR motif and model perplexity were analyzed. In natural Tat signal peptides, the position of the first RR motif gradually shifted toward the C-terminal direction as sequence length increased, whereas RR motifs in ApexSP-generated sequences were predominantly maintained within the first one-third of the sequence (Fig. 2l), indicating that the model retained the positional constraint of the N-terminal twin-arginine motif required for Tat pathway recognition. Meanwhile, generated sequences maintained low perplexity across different length ranges (Sec: 2.9–4.2; Tat: 2.4–3.4; Fig. 2m), suggesting stable amino acid recovery capability across diverse sequence lengths. Collectively, these analyses demonstrate that ApexSP-generated sequences exhibit distinct generation preferences characteristic of secretion signal peptides.

### 3.3 ApexSP enables robust joint prediction of signal peptide pathway, topology, and taxonomic origin

To constrain generated sequences that may deviate from the biological requirements of authentic secretion signals, ApexSP incorporates a biological filter designed to simultaneously predict signal peptide pathway, residue-level topology regions, and species origin, while accommodating different input formats, including full-length signal peptides, N-terminal fragments of secreted proteins, and truncated sequences (Fig. 3a).

**Figure 3.**
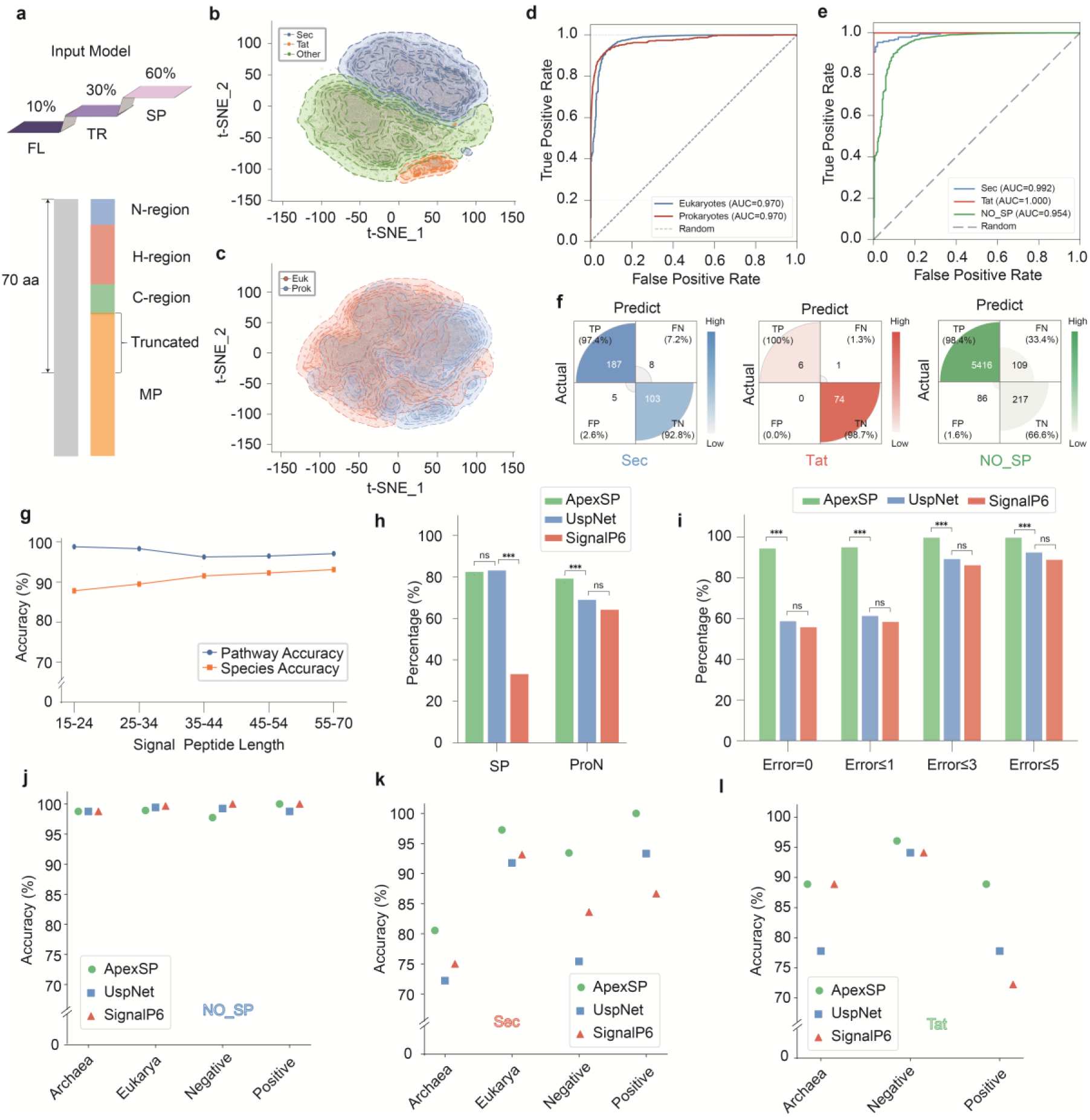
ApexSP enables joint prediction of signal peptide pathways, topology, and species attributes. **(a)** Schematic representation of the input strategy for the topology-aware multitask predictor in ApexSP. **(b)** t-SNE visualization of Sec, Tat, and Other signal peptides in the training set. **(c)** t-SNE visualization of eukaryotic and prokaryotic signal peptides in the training set. **(d)** ROC curve of ApexSP for species-origin classification on the Signal6_Set validation set. **(e)** ROC curve of ApexSP for pathway classification of Sec, Tat, and Other classes on the Signal6_Set validation set. **(f)** Confusion matrices of ApexSP for Sec, Tat, and Other classification on the Signal6_Set validation set. **(g)** Pathway and species-origin classification accuracy of ApexSP across different signal peptide length groups. **(h)** Comparison of pathway classification accuracy among ApexSP, USPNet, and SignalP6 on the SPSDB dataset using full signal peptides (SP) and protein N-terminal fragments (ProN) as inputs. **(i)** Comparison of cleavage-site prediction accuracy among ApexSP, USPNet, and SignalP6 under different positional error thresholds on the SPSDB dataset. **(j)** Comparison of classification accuracy among ApexSP, USPNet, and SignalP6 for non-signal peptide (NO_SP) sequences on the Usp_bench dataset. **(k)** Comparison of classification accuracy among ApexSP, USPNet, and SignalP6 for Sec signal peptides on the Usp_bench dataset. **(l)** Comparison of classification accuracy among ApexSP, USPNet, and SignalP6 for Tat signal peptides on the Usp_bench dataset.

In the learned representation space, Sec, Tat, and No_SP sequences showed largely separable distributions, with Tat sequences occupying a relatively distinct region and a visible boundary also emerging between Sec and No_SP sequences. (Fig. 3b). By contrast, the representations of eukaryotic and prokaryotic sequences showed greater overlap, although partial separation was still observed (Fig. 3c). Ablation analysis showed that incorporation of ESM2 pretrained representations increased taxonomic-origin classification accuracy from 0.9297 to 0.9622. Further introduction of residue-level topology supervision largely preserved classification performance and enabled cleavage-boundary prediction. (Table S8).

On the external Signal6_Set validation dataset, ApexSP achieved ROC–AUC values of 0.970 for both eukaryotic and prokaryotic origin classification (Fig. 3d). For pathway classification, the ROC–AUC values for Sec, Tat, and NO_SP were 0.992, 1.000, and 0.954, respectively (Fig. 3e). The confusion matrices showed that precision and recall were 97.4% and 95.9% for Sec, 100% and 85.7% for Tat, and 98.4% and 98.0% for NO_SP, respectively (Fig. 3f). These results show that ApexSP distinguished all three pathway classes with high precision, although recall was comparatively lower for the smaller Tat subset.

Model performance also remained stable across different signal peptide length ranges. In the independent UniRef50 validation set, pathway classification accuracy was no lower than 96.2% across all length groups from 15 to 70 aa, whereas taxonomic-origin classification accuracy gradually increased from 87.8% in the shortest sequence group to 93.1% in the longest group (Fig. 3g). Thus, species accuracy increased with sequence length, whereas pathway classification remained comparatively insensitive to input length.

Robustness of pathway prediction under complete signal peptide and secreted-protein N-terminal fragment inputs was subsequently compared on the challenging SPSDB dataset, which contains diverse signal peptides and non-secretory proteins. When isolated signal peptide sequences were used as input, professional SignalP 6 frequently classified Sec signal peptides as non-secretory, resulting in an accuracy of 33.1%., whereas ApexSP achieved 82.4%, close to the 83.1% obtained by USPNet. When secreted-protein N-terminal fragments were used as input, ApexSP retained an accuracy of 79.2%, substantially higher than the 68.9% and 64.2% achieved by USPNet and SignalP 6, respectively (Fig. 3h). These results show that ApexSP maintained robust performance across different sequence input types.

For cleavage-site localization, ApexSP achieved an accuracy of 94.3% under the strict exact-match criterion, significantly higher than 58.6% for USPNet and 55.6% for SignalP 6. When an error of one residue was allowed, the accuracies of the three methods were 94.9%, 61.2%, and 58.3%, respectively. When an error of three residues was allowed, ApexSP reached 99.6% accuracy (Fig. 3i). These results indicate that ApexSP resolved the cleavage boundary between the signal peptide and mature protein more precisely, with the largest advantage observed under strict cleavage-site prediction criteria.

Finally, ApexSP, USPNet, and SignalP 6 were systematically compared across taxonomic subsets of the SP22 dataset. In the NO_SP task, all three methods achieved accuracies above 99% in the archaeal, eukaryotic, Gram-negative, and Gram-positive subsets (Fig. 3j). For Sec signal peptides, ApexSP achieved accuracies of 80.6%, 97.3%, 93.4%, and nearly 100% in the archaeal, eukaryotic, Gram-negative, and Gram-positive subsets, respectively, and generally outperformed USPNet and SignalP 6 (Fig. 3k). For Tat signal peptides, ApexSP achieved accuracies of approximately 88.9%, 96.1%, and 88.9% in the archaeal, Gram-negative, and Gram-positive subsets, respectively, again outperforming USPNet and SignalP 6 (Fig. 3l). Collectively, ApexSP maintained robust multi-task performance across different sequence inputs, sequence lengths, and taxonomic groups, providing a reliable foundation for the biological constraint filtering of generated signal peptides.

### 3.4 ApexSP enables cargo-aware prioritization of signal peptides

To determine the effective contribution range of mature protein sequence information for signal peptide selection, we first evaluated the impact of different lengths of N-terminal mature protein fragments on SP–MP compatibility prediction. The random baseline yielded a Spearman correlation close to 0, whereas all models incorporating mature-protein N-terminal sequences showed positive correlations. Spearman correlations for the 10-, 50-, 100-, and 200-aa inputs were mainly distributed between 0.15 and 0.25, with the 200-aa configuration showing relatively high overall performance and lower variation across data splits. Extending the input length to 500 aa did not further improve ranking performance (Fig. 4a). Therefore, the N-terminal 200 residues of the mature protein were used as input for subsequent compatibility modeling.

**Figure 4.**
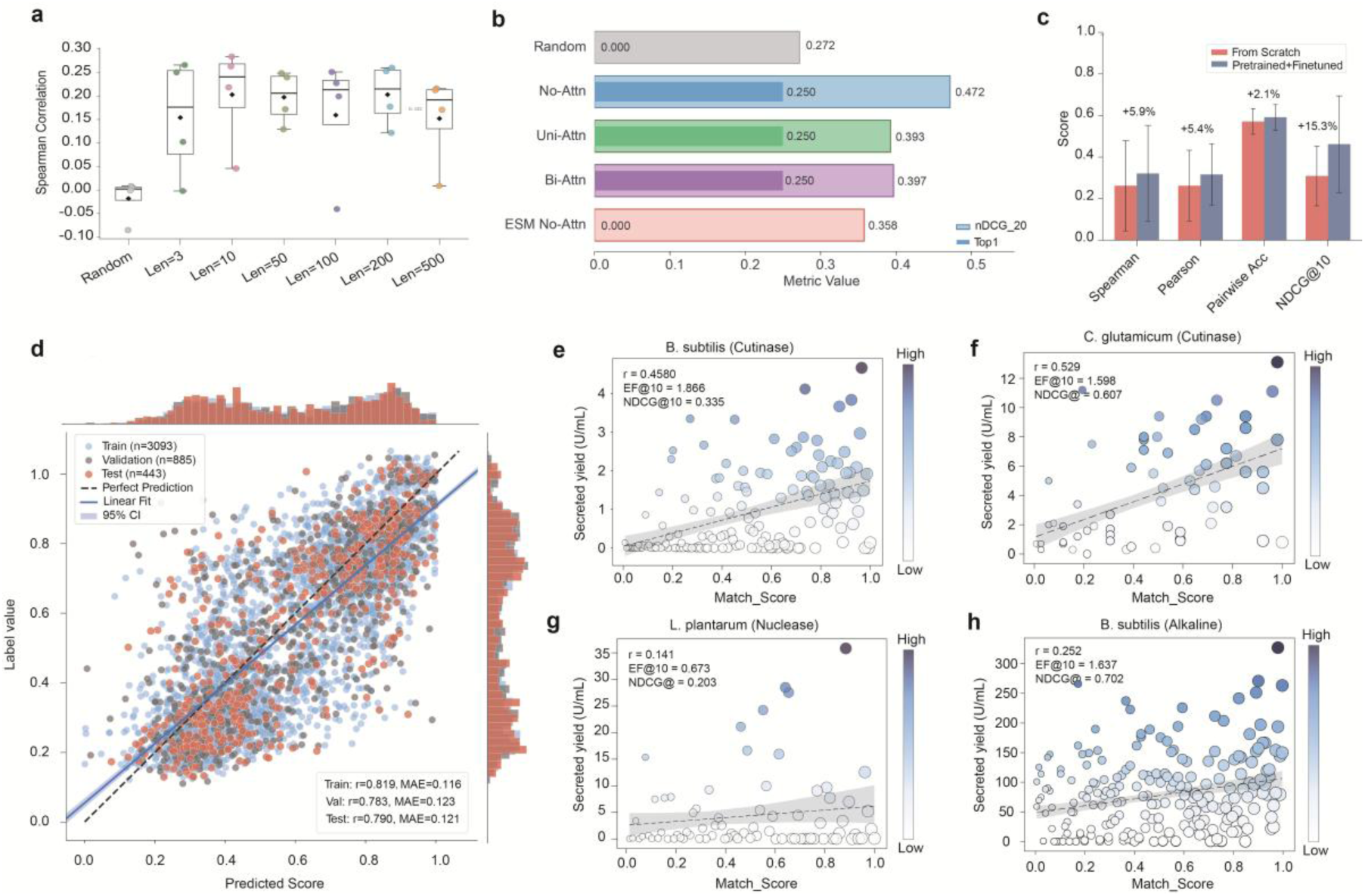
ApexSP protein-aware ranking model predicts the secretion potential of signal peptide–mature protein combinations. **(a)** Effect of mature protein N-terminal input length on SP–MP compatibility ranking performance. **(b)** Structural ablation analysis of the SP–MP compatibility model. **(c)** Comparison of models trained from scratch and pretrained models followed by fine-tuning. **(d)** Prediction performance of the fine-tuned ApexSP model in the AmyQ– Bacillus subtilis secretion system. **(e)** Secretion ranking validation of ApexSP in the Bacillus subtilis–cutinase system. r denotes the correlation coefficient, EF@10 represents enrichment of high-yield candidates among the top 10% ranked candidates, and NDCG@10 evaluates the ranking quality of top candidates. **(f)** Secretion ranking validation of ApexSP in the Corynebacterium glutamicum–cutinase system. **(g)** Secretion ranking validation of ApexSP in the Lactiplantibacillus plantarum WCFS1–nuclease system. **(h)** Secretion ranking validation of ApexSP in the Bacillus subtilis–alkaline xylanase system.

Subsequently, we compared the effects of different sequence representations and interaction architectures on ranking performance. The results showed that the dual-tower model without explicit cross-attention achieved the best ranking performance (nDCG@20 = 0.472, Top-1 = 0.250), whereas introducing unidirectional or bidirectional cross-attention did not further improve ranking capability (Fig. 4b). In addition, incorporating representations from the general protein language model ESM2 provided no additional benefit, suggesting that task-specific modeling of SP–MP compatibility relationships is more critical than generic sequence representation for this task^53^.

Pseudo-label pretraining further improved ranking performance on experimental secretion data. Compared with a randomly initialized model trained from scratch, the pretrained and fine-tuned model showed improvements of 5.9%, 5.4%, 2.1%, and 15.3% in Spearman correlation, Pearson correlation, pairwise accuracy, and NDCG@10, respectively, with the largest improvement observed for NDCG@10 (Fig. 4c). In the AmyQ–*Bacillus subtilis* system from the SPE dataset, the model prediction scores showed consistent correlation trends with normalized secretion yields across the training, validation, and test sets, with Pearson’s *r* values of 0.819, 0.783, and 0.790, respectively (Fig. 4d). The comparable performance between the validation and test sets indicates that the model can learn SP–MP compatibility patterns from limited experimental data.

Next, we evaluated the cross-system ranking capability of ApexSP in four independent expression systems. In the *B. subtilis*–cutinase expression system, the predicted ranking showed a positive correlation with experimental secretion yields (*r* = 0.458), with a 1.866-fold enrichment of high-yield candidates within the top 10% predictions (Fig. 4e). In the *C. glutamicum*–cutinase system, the model achieved stronger correlation (*r* = 0.529) and improved ranking performance (NDCG@10 = 0.607) (Fig. 4f). In the *L. plantarum* nuclease expression system, the overall prediction correlation was weaker (*r* = 0.141) (Fig. 4g). However, in the *B. subtilis*–alkaline xylanase system, the model still enriched high-yield signal peptides among the top-ranked candidates (EF@10 = 1.637, NDCG@10 = 0.702) (Fig. 4h). Collectively, these results across diverse expression systems demonstrate that ApexSP can leverage cargo protein context information to prioritize signal peptide candidates with higher secretion potential.

### 3.5 ApexSP-designed signal peptides outperform benchmark signal peptides in secretion performance

To evaluate the actual secretion performance of ApexSP-designed signal peptides, we experimentally tested them across different hosts and secretion pathways, including the *Pichia pastoris* ApGA expression system and the Sec- and Tat-dependent secretion systems of *Bacillus subtilis* expressing AprE2.

After completing the ApexSP design process, we first examined the predicted performance of the designed signal peptides. In the ApGA–*P. pastoris* system, the compatibility scores of ApexSP candidates were mainly distributed within the range of 0.35–0.55 (Fig. 5a). In the AprE2–*B. subtilis* system, both Sec and Tat candidates exhibited high predicted compatibility scores, with mean values of 0.55 and 0.65, respectively, and Tat signal peptides showed overall higher scores than Sec candidates (Fig. 5b). Based on the SP–MP compatibility model predictions, the top-ranked 10 candidate signal peptides from each system were selected for subsequent experimental validation. We next compared the predicted scores between ApexSP candidates and reference signal peptides. In the ApGA–*P. pastoris* system, the ten candidate signal peptides showed slightly higher predicted scores than the α-factor reference signal peptide, with scores concentrated around 0.55 compared with approximately 0.4 for α-factor (Fig. 5c). In the AprE2–*B. subtilis* Sec system, ApexSP-designed candidates exhibited comparable prediction scores to the Q99405-derived reference signal peptide, with both distributed at a high score level of approximately 0.75. In contrast, in the AprE2–*B. subtilis* Tat system, the commonly used PhoD signal peptide showed a relatively low predicted score (∼0.3), whereas the designed Tat signal peptides achieved substantially higher scores (∼0.8) (Fig. 5d).

**Figure 5.**
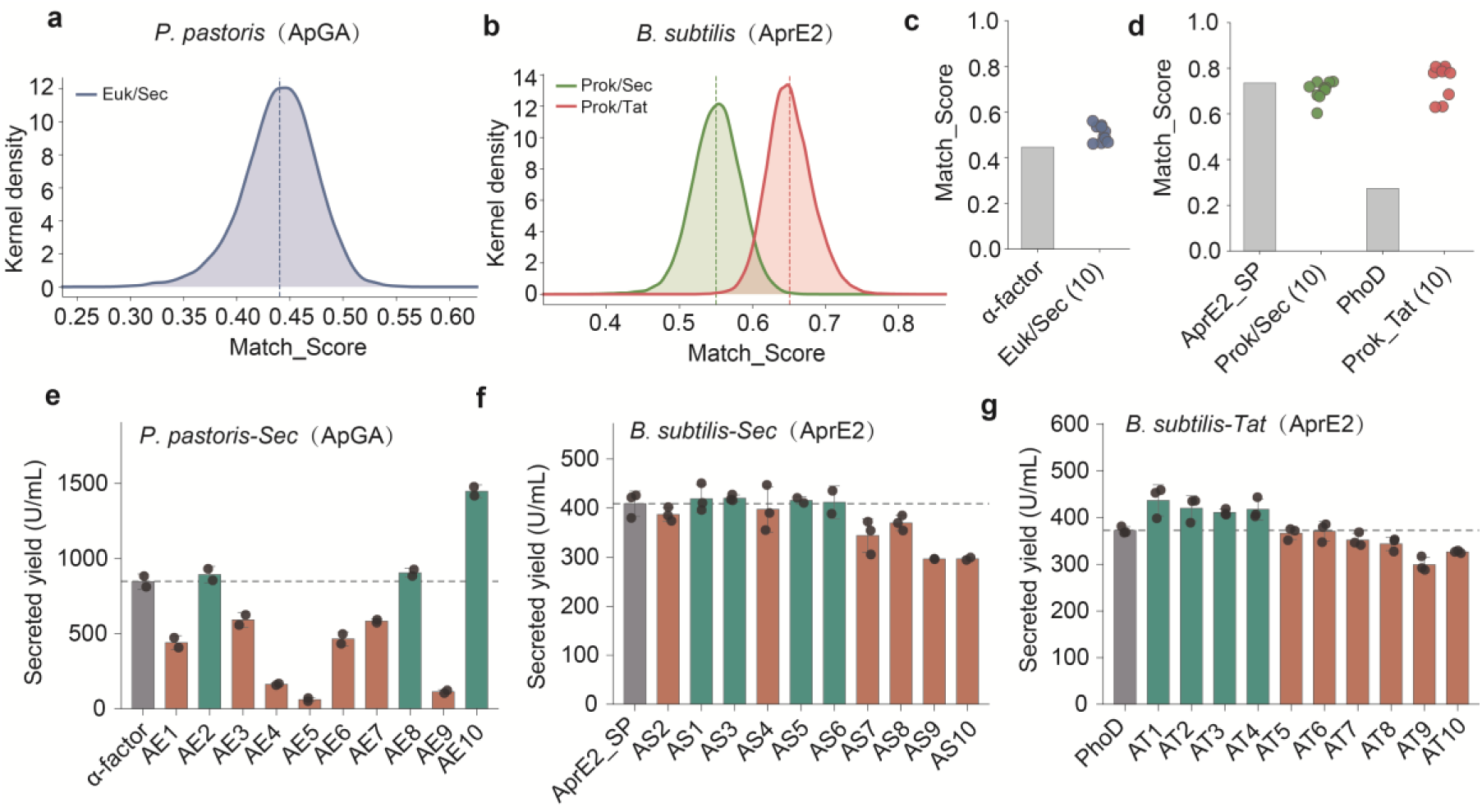
Experimental validation of ApexSP-designed signal peptides in heterologous secretion systems. **(a)** Predicted compatibility score distribution of ApexSP-designed signal peptides for ApGA secretion in *Pichia pastoris*. **(b)** Predicted compatibility score distribution of ApexSP-designed signal peptides for AprE2 secretion through Sec and Tat pathways in *Bacillus subtilis*. **(c)** Comparison of predicted compatibility scores between the α-factor signal peptide and ApexSP-designed candidates for ApGA secretion in *P. pastoris*. **(d)** Comparison of predicted compatibility scores between benchmark signal peptides and ApexSP-designed candidates for AprE2 secretion through Sec and Tat pathways in *B. subtilis*. **(e)** Secreted glucoamylase activity of ApGA fused with the α-factor signal peptide or ApexSP-designed signal peptides in *P. pastoris*. **(f)** Secreted protease activity of AprE2 fused with the AprE2-derived benchmark signal peptide or ApexSP-designed Sec signal peptides in *B. subtilis*. **(g)** Secreted protease activity of AprE2 fused with the PhoD signal peptide or ApexSP-designed Tat signal peptides in *B. subtilis*.

Experimental results further demonstrated the ability of the model ranking to enrich high-performance signal peptides. In the ApGA–*P. pastoris* system, the α-factor signal peptide produced an extracellular enzyme activity of 847.5 U/mL, whereas the ApexSP-designed signal peptide AE10 achieved 1,448.3 U/mL, representing an approximately 1.71-fold improvement over the reference signal peptide. In addition, AE2 and AE8 also exceeded the secretion performance of α-factor (Fig. 5e). In the AprE2–*B. subtilis* Sec system, all ten designed signal peptides maintained high secretion levels, with four candidates exceeding the endogenous AprE2 reference signal peptide (409.1 U/mL) (Fig. 5f).

In the AprE2–*B. subtilis* Tat secretion system, the 10 high-ranked candidate signal peptides also maintained high secretion performance, with all candidates achieving extracellular protease activities above 300 U/mL. The PhoD reference signal peptide resulted in an extracellular protease activity of 372.4 U/mL, whereas four designed signal peptides exceeded 410 U/mL. Among them, AT1 showed the highest secretion performance, reaching an extracellular protease activity of 436.7 U/mL (Fig. 5g). In summary, ApexSP enables the enrichment of high-performance signal peptide candidates that outperform commonly used reference signal peptides across diverse cargo proteins, hosts, and secretion pathways.

## 4. Discussion

Protein secretion performance is a complex trait jointly determined by signal peptide sequence, mature protein properties, secretion pathway selection, and host background^9–11^. ApexSP represents an important step toward addressing this challenge by incorporating these contextual factors into a unified design framework. Through conditional generation, biological filtering, and cargo protein-aware ranking, ApexSP not only generates Sec and Tat candidate signal peptides that follow natural signal peptide characteristics, but also evaluates their compatibility with specific cargo proteins. Experimental validation demonstrated that signal peptides designed by ApexSP achieved directly applicable high-secretion performance across multiple expression systems, outperforming commonly used reference signal peptides.

An important foundation of this study is the construction of a large-scale, high-confidence signal peptide database. Although signal peptide sequences are abundant in public databases, existing resources are typically annotated for individual attributes and lack a unified representation integrating secretion pathway, topology boundaries, and taxonomic origin information^54^. By integrating UniRef50, UniProtKB/Swiss-Prot, and publicly available validation datasets, together with consistent annotation from SignalP6 and USPNet, ApexSP associates signal peptide sequences with secretion pathways, topological regions, and NCBI Taxonomy-derived source information, thereby establishing a signal peptide knowledge base with multidimensional attributes. This strategy partially reduces annotation bias caused by reliance on a single prediction tool and, more importantly, provides additional information regarding taxonomic origin and cargo protein associations. Based on this knowledge system, ApexSP enables a transition from learning natural signal peptide patterns to designing signal peptides under host- and cargo protein-specific constraints.

Second, a major contribution of ApexSP is that it advances signal peptide design toward experimentally actionable performance. Most existing signal peptide generation models remain focused on confirming whether generated sequences possess secretion-related characteristics^15,16^. However, designed signal peptides only achieve clear engineering value when they can efficiently direct secretion of specific cargo proteins and outperform commonly used or universal signal peptides experimentally. In ApGA–*P. pastoris* and AprE2–*B. subtilis* systems, multiple candidates among only 10 experimentally tested signal peptides designed by ApexSP exceeded the performance of highly effective reference signal peptides. Particularly in the ApGA–*P. pastoris* system, candidate signal peptides with higher compatibility scores consistently achieved extracellular enzyme activities above 300 U/mL. These results demonstrate that ApexSP can effectively predict and enrich candidate signal peptides with high secretion potential.

The overall ApexSP framework addresses three key challenges in signal peptide design through a hierarchical strategy. The generation module learns the sequence principles of natural Sec and Tat signal peptides from large-scale datasets and expands the candidate space available for screening. The biological filtering module further constrains candidate sequences according to secretion pathway, topological boundaries, and species origin, thereby excluding sequences that do not conform to fundamental secretion-associated rules. The cargo protein-aware ranking module then leverages signal peptide–mature protein pairing relationships to prioritize candidates according to their potential to drive the secretion of specific proteins. Through this generation– constraint–ranking strategy, ApexSP connects native-like signal peptide sequences with cargo protein-oriented functional selection. In the future, secretion potential and biological constraints could be integrated directly into the generation process^55^, enabling simultaneous optimization of biological validity and cargo protein compatibility during sequence generation and thereby producing signal peptides that are more precisely adapted to specific expression systems.

In summary, this study establishes a cargo protein–customized signal peptide engineering framework, advancing signal peptide research from conventional recognition and screening approaches toward target-constrained active design. With the accumulation of more standardized secretion datasets and the integration of host-specific information, ApexSP has the potential to further evolve into an intelligent secretion element design platform for complex protein expression systems.

## Acknowledgments

This work was supported by the National Key Research and Development Program of China (2023YFC3403502), National Natural Science Foundation of China (32301041, 32571437, 32401034), SKLMT Frontier and Challenges Project (SKLMTFCP-2023-01), SKLDRS Open Project (2025SKLDRS0323), and Intramural Joint Program Fund of SKLMT (SKLMTIJP-2025-03). X.J. is supported by TaiShan Scholars (NO.tsqn202507080).

## Author contributions

Chunyi Yang, Ruiyang Hou, Zhenyu Ma, Qiuyan Kang, ShuoJing Zuo, Limei Xu: Data curation; formal analysis; investigation; methodology; writing – original draft; writing – review & editing. Min Xiao: Supervision; resources. Xiuyun Wu, Xukai Jiang: Conceptualization; supervision; project administration; funding acquisition; writing – review & editing.

## Data availability statement

The ApexSP inference code and model weights are publicly available at https://huggingface.co/ycy77/ApexSP.

## Ethics statement

No animals or humans were involved in this study.

## Additional Information

Supplementary information includes:

**Table S1.** Strains used in this study.

**Table S2.** Codon-optimized DNA sequences used for gene synthesis.

**Table S3.** Amino acid sequences of signal peptides used for expression optimization experiments.

**Table S4.** DNA primers used for signal peptide replacement.

**Table S5.** Signal peptide dataset compiled in this study.

**Table S6.** Assessment of sequence overlap between training and external validation datasets.

**Table S7.** Ablation analysis of the evolution-guided discrete diffusion model.

**Table S8.** Ablation analysis of the topology-aware multitask biological filtering model.

