## Supporting_Information for "Beyond Generic Signal Peptides: ApexSP Enables Cargo-Specific Secretion Design"

**Running title:** Beyond Universal Signal Peptides

**This file contains:**

**Table S1.** Strains used in this study.

**Table S2.** Codon-optimized DNA sequences used for gene synthesis.

**Table S3.** Amino acid sequences of signal peptides used in expression optimization experiments.

**Table S4.** DNA primers used in replacement of signal peptides.

**Table S5.** The signal peptide dataset compiled in this study.

**Table S6.** Dataset Overlap Assessment.

**Table S7.** Ablation analysis of the evolution-guided discrete diffusion model.

**Table S8.** Ablation analysis of the topology-aware multitask biological filter.

**Figure S1.** The probability density plot of transmembrane free energy barriers in the BDBS dataset.

**Tables**

**Table S1.** Strains used in this study

| **Strains names** | **Characteristics** | **Reference** |
|---|---|---|
| DH5α | *Escherichia coli* | Vazyme |
| X-33 | *Pichia pastoris* | Lab stock |
| WB600 | *Bacillus subtilis* | Lab stock |

**Table S2.** DNA sequences of proteins synthesized after codon optimization

| Names | Sequence (5'-3') |
|---|---|
| ApGA | GCTACTGGTTCTCTGTCTAGTTGGTTGAGTTCCGAGAATACTGTTG<br>CTTTGCAAGGTGTATTGAACAACATCGGAGCTAGTGGTTCTAAGGC<br>TTCTGGTGCTAGTGCAGGTGTTGTTGTTGCTAGTCCATCTAAGTCTA<br>ATCCAGACTACTTCTACACTTGGACCAGAGATTCTGCCTTGGTCTTT<br>AAGGCTCTAGTTGACCAGTTGATCGCCGGTAATAAGAGTTTGGAAC<br>CACTTATCCAGCAATACATCTCTGCTCAGGCTAAGTTGCAAACTGTT<br>AACAATCCATCCGGTGGTCTGTGTTCTGGTGGTTTGGCTGAACCTA<br>AGTTTGAGGTTGATCTGACTCCATTTACTGGTGCTTGGGGTAGACC<br>TCAAAGAGATGGACCAGCTTTGAGAGCTACCGCTATGATCGCTTAC<br>AGTAGATACTTGATCGCTAACGGTAACACCACCACTGTCAACAACAT<br>CATCTGGCCTATCGTCCAGAATGACCTGTCTTATGTCACTCAATACT<br>GGAATCAAACTACCTTTGATCTGTGGGAAGAAATCAACTCTTCATCT<br>TTCTTCACTACCGCAGTTCAGTACAGAGCTTTGGTTGAGGGTAACA<br>ACTTGGCTACTCAATTGGGTAAGAGTTGTCCAAACTGTGTATCACAA<br>GCTCCATTGGTCTTGTGCTTTCTTCAGAGTTACTGGACTGGATCTTA<br>CGCTTTGTCAAATACTGGTGGAGGTAGATCTGGTAAGGACGCTAAC<br>AGTATTCTGACCTCCATCCATATCTTTGACCCTGCAGCTTCTTGTGA<br>CAGTACCACCTTTCAACCTTGTTCCGACAAGGCTTTGGCAAATCATA<br>AGGTTGTTACCGATTCTTTCAGATCCATCTACTCCATTAATCAGGGT<br>ATCGCTCAAGGTTCCGGAGTTGCTGTTGGAAGATATCCAGAGGACT<br>CTTACTACAATGGTAATCCATGGTATTTGAACACTTTCGCTGCTGCT<br>GAACAATTGTACGATGCTGTTTACCAGTGGAAGAAGATTGGTTCTAT<br>CTCCATCACATCCATTTCTTTGCCATTCTTCAAAGATGTCTACTCTAG<br>TGCTGCTGTTGGCACCTATTCATCCAGTACCGTTACCTTCACCTCTA<br>TCGTTAATGCTGTCCAGACTTATGCAGACTCTTACATGTCCATCGCT<br>CAGAAGTACACACCTTCTAATGGTGCTTTGTCTGAACAATACAACAG<br>AGCTGACGGTACTCCATTGTCTGCTGTCGATCTGACCTGGAGTTAT<br>GCTGCCTTCTTGACTGCTTACAATGCAAGAGCAAACGTTCTGCCAG<br>CATCTTGGGGAGCTAGTTCTGCTAAGCTGCCTAACTCTTGTTCATC<br>CGGATCTGCAACTGGTCCATGTGCTGCAGCTACCAACACTAACTGG<br>GGTAATCCAGGCTCTCCAAGTACTGGAACTCCAACTACCACTACTG<br>GTGGATCT |
| pPICZA-ApGA | AGATCTAACATCCAAAGACGAAAGGTTGAATGAAACCTTTTTGCCAT<br>CCGACATCCACAGGTCCATTCTCACACATAAGTGCCAAACGCAACA<br>GGAGGGGATACACTAGCAGCAGACCGTTGCAAACGCAGGACCTCC<br>ACTCCTCTTCTCCTCAACACCCACTTTTGCCATCGAAAAACCAGCC<br>CAGTTATTGGGCTTGATTGGAGCTCGCTCATTCCAATTCCTTCTATT<br>AGGCTACTAACACCATGACTTTATTAGCCTGTCTATCCTGGCCCCCC<br>TGGCGAGGTTCATGTTTGTTTATTTCCGAATGCAACAAGCTCCGCAT<br>TACACCCGAACATCACTCCAGATGAGGGCTTTCTGAGTGTGGGGT<br>CAAATAGTTTCATGTTCCCCAAATGGCCCAAAACTGACAGTTTAAAC<br>GCTGTCTTGGAACCTAATATGACAAAAGCGTGATCTCATCCAAGATG |

AACTAAGTTTGGTTCGTTGAAATGCTAACGGCCAGTTGGTCAAAAA
GAAACTTCCAAAAGTCGGCATACCGTTTGTCTTGTTTGGTATTGATT
GACGAATGCTCAAAAATAATCTCATTAATGCTTAGCGCAGTCTCTCT
ATCGCTTCTGAACCCCGGTGCACCTGTGCCGAAACGCAAATGGGG
AAACACCCGCTTTTTGGATGATTATGCATTGTCTCCACATTGTATGC
TTCCAAGATTCTGGTGGGAATACTGCTGATAGCCTAACGTTCATGAT
CAAAATTTAACTGTTCTAACCCCTACTTGACAGCAATATATAAACAGA
AGGAAGCTGCCCTGTCTTAAACCTTTTTTTTTATCATCATTATTAGCT
TACTTTCATAATTGCGACTGGTTCCAATTGACAAGCTTTTGATTTTAA
CGACTTTTAACGACAACTTGAGAAGATCAAAAAACAACTAATTATTC
GAAACGATGAGATTTCCTTCAATTTTTACTGCTGTTTTATTCGCAGC
ATCCTCCGCATTAGCTGCTCCAGTCAACACTACAACAGAAGATGAA
ACGGCACAAATTCCGGCTGAAGCTGTCATCGGTTACTCAGATTTAG
AAGGGGATTTCGATGTTGCTGTTTTGCCATTTTCCAACAGCACAAAT
AACGGGTTATTGTTTATAAATACTACTATTGCCAGCATTGCTGCTAAA
GAAGAAGGGGTATCTCTCGAGAAAAGAGCTACTGGTTCTCTGTCTA
GTTGGTTGAGTTCCGAGAATACTGTTGCTTTGCAAGGTGTATTGAA
CAACATCGGAGCTAGTGGTTCTAAGGCTTCTGGTGCTAGTGCAGGT
GTTGTTGTTGCTAGTCCATCTAAGTCTAATCCAGACTACTTCTACAC
TTGGACCAGAGATTCTGCCTTGGTCTTTAAGGCTCTAGTTGACCAG
TTGATCGCCGGTAATAAGAGTTTGGAACCACTTATCCAGCAATACAT
CTCTGCTCAGGCTAAGTTGCAAACTGTTAACAATCCATCCGGTGGT
CTGTGTTCTGGTGGTTTGGCTGAACCTAAGTTTGAGGTTGATCTGA
CTCCATTTACTGGTGCTTGGGGTAGACCTCAAAGAGATGGACCAGC
TTTGAGAGCTACCGCTATGATCGCTTACAGTAGATACTTGATCGCTA
ACGGTAACACCACCACTGTCAACAACATCATCTGGCCTATCGTCCA
GAATGACCTGTCTTATGTCACTCAATACTGGAATCAAACTACCTTTG
ATCTGTGGGAAGAAATCAACTCTTCATCTTTCTTCACTACCGCAGTT
CAGTACAGAGCTTTGGTTGAGGGTAACAACTTGGCTACTCAATTGG
GTAAGAGTTGTCCAAACTGTGTATCACAAGCTCCATTGGTCTTGTG
CTTTCTTCAGAGTTACTGGACTGGATCTTACGCTTTGTCAAATACTG
GTGGAGGTAGATCTGGTAAGGACGCTAACAGTATTCTGACCTCCAT
CCATATCTTTGACCCTGCAGCTTCTTGTGACAGTACCACCTTTCAAC
CTTGTTCCGACAAGGCTTTGGCAAATCATAAGGTTGTTACCGATTCT
TTCAGATCCATCTACTCCATTAATCAGGGTATCGCTCAAGGTTCCGG
AGTTGCTGTTGGAAGATATCCAGAGGACTCTTACTACAATGGTAATC
CATGGTATTTGAACACTTTCGCTGCTGCTGAACAATTGTACGATGCT
GTTTACCAGTGGAAGAAGATTGGTTCTATCTCCATCACATCCATTTC
TTTGCCATTCTTCAAAGATGTCTACTCTAGTGCTGCTGTTGGCACCT
ATTCATCCAGTACCGTTACCTTCACCTCTATCGTTAATGCTGTCCAG
ACTTATGCAGACTCTTACATGTCCATCGCTCAGAAGTACACACCTTC
TAATGGTGCTTTGTCTGAACAATACAACAGAGCTGACGGTACTCCAT
TGTCTGCTGTCGATCTGACCTGGAGTTATGCTGCCTTCTTGACTGC
TTACAATGCAAGAGCAAACGTTCTGCCAGCATCTTGGGGAGCTAGT

TCTGCTAAGCTGCCTAACTCTTGTTCATCCGGATCTGCAACTGGTC
CATGTGCTGCAGCTACCAACACTAACTGGGGTAATCCAGGCTCTCC
AAGTACTGGAACTCCAACTACCACTACTGGTGGATCTCATCATCATC
ATCATCATTGAGTTTGTAGCCTTAGACATGACTGTTCCTCAGTTCAA
GTTGGGCACTTACGAGAAGACCGGTCTTGCTAGATTCTAATCAAGA
GGATGTCAGAATGCCATTTGCCTGAGAGATGCAGGCTTCATTTTTG
ATACTTTTTTATTTGTAACCTATATAGTATAGGATTTTTTTTGTCATTTT
GTTTCTTCTCGTACGAGCTTGCTCCTGATCAGCCTATCTCGCAGCT
GATGAATATCTTGTGGTAGGGGTTTGGGAAAATCATTCGAGTTTGAT
GTTTTTCTTGGTATTTCCCACTCCTCTTCAGAGTACAGAAGATTAAG
TGAGACCTTCGTTTGTGCGGATCCCCCACACACCATAGCTTCAAAA
TGTTTCTACTCCTTTTTTACTCTTCCAGATTTTCTCGGACTCCGCGC
ATCGCCGTACCACTTCAAAACACCCAAGCACAGCATACTAAATTTCC
CCTCTTTCTTCCTCTAGGGTGTCGTTAATTACCCGTACTAAAGGTTT
GGAAAAGAAAAAAGAGACCGCCTCGTTTCTTTTTCTTCGTCGAAAA
AGGCAATAAAAATTTTTATCACGTTTCTTTTTCTTGAAAATTTTTTTT
TTGATTTTTTTCTCTTTCGATGACCTCCCATTGATATTTAAGTTAATAA
ACGGTCTTCAATTTCTCAAGTTTCAGTTTCATTTTTCTTGTTCTATTA
CAACTTTTTTTACTTCTTGCTCATTAGAAAGAAAGCATAGCAATCTAA
TCTAAGGGCGGTGTTGACAATTAATCATCGGCATAGTATATCGGCAT
AGTATAATACGACAAGGTGAGGAACTAAACCATGGCCAAGTTGACC
AGTGCCGTTCCGGTGCTCACCGCGCGCGACGTCGCCGGAGCGGT
CGAGTTCTGGACCGACCGGCTCGGGTTCTCCCGGGACTTCGTGG
AGGACGACTTCGCCGGTGTGGTCCGGGACGACGTGACCCTGTTC
ATCAGCGCGGTCCAGGACCAGGTGGTGCCGGACAACACCCTGGC
CTGGGTGTGGGTGCGCGGCCTGGACGAGCTGTACGCCGAGTGGT
CGGAGGTCGTGTCCACGAACTTCCGGGACGCCTCCGGGCCGGCC
ATGACCGAGATCGGCGAGCAGCCGTGGGGGCGGGAGTTCGCCCT
GCGCGACCCGGCCGGCAACTGCGTGCACTTCGTGGCCGAGGAGC
AGGACTGACACGTCCGACGCGGCCCGACGGGTCCGAGGCCTCG
GAGATCCGTCCCCCTTTTCCTTTGTCGATATCATGTAATTAGTTATGT
CACGCTTACATTCACGCCCTCCCCCCACATCCGCTCTAACCGAAAA
GGAAGGAGTTAGACAACCTGAAGTCTAGGTCCCTATTTATTTTTTTA
TAGTTATGTTAGTATTAAGAACGTTATTTATATTTCAAATTTTTCTTTTT
TTTCTGTACAGACGCGTGTACGCATGTAACATTATACTGAAAACCTT
GCTTGAGAAGGTTTTGGGACGCTCGAAGGCTTTAATTTGCAAGCTG
GAGACCAACATGTGAGCAAAAGGCCAGCAAAAGGCCAGGAACCGT
AAAAAGGCCGCGTTGCTGGCGTTTTTCCATAGGCTCCGCCCCCCT
GACGAGCATCACAAAAATCGACGCTCAAGTCAGAGGTGGCGAAAC
CCGACAGGACTATAAAGATACCAGGCGTTTCCCCCTGGAAGCTCCC
TCGTGCGCTCTCCTGTTCCGACCCTGCCGCTTACCGGATACCTGT
CCGCCTTTCTCCCTTCGGGAAGCGTGGCGCTTTCTCATAGCTCAC
GCTGTAGGTATCTCAGTTCGGTGTAGGTCGTTCGCTCCAAGCTGG
GCTGTGTGCACGAACCCCCCGTTCAGCCCGACCGCTGCGCCTTAT

| | |
|---|---|
| | CCGGTAACTATCGTCTTGAGTCCAACCCGGTAAGACACGACTTATC<br>GCCACTGGCAGCAGCCACTGGTAACAGGATTAGCAGAGCGAGGTA<br>TGTAGGCGGTGCTACAGAGTTCTTGAAGTGGTGGCCTAACTACGG<br>CTACACTAGAAGAACAGTATTTGGTATCTGCGCTCTGCTGAAGCCA<br>GTTACCTTCGGAAAAAGAGTTGGTAGCTCTTGATCCGGCAAACAAA<br>CCACCGCTGGTAGCGGTGGTTTTTTTGTTTGCAAGCAGCAGATTAC<br>GCGCAGAAAAAAAGGATCTCAAGAAGATCCTTTGATCTTTTCTACG<br>GGGTCTGACGCTCAGTGGAACGAAAACTCACGTTAAGGGATTTTG<br>GTCATGAGATC |
| aprE2 | GCTGAAGAAGCAAAAGAAAAATATTTAATTGGCTTTAATGAGCAGGA<br>AGCTGTCAGTGAGTTTGTAGAACAAGTAGAGGCAAATGACGAGGT<br>CGCCATTCTCTCTGAGGAAGAGGAAGTCGAAATTGAATTGCTTCAT<br>GAATTTGAAACGATTCCTGTTTTATCCGTTGAGTTAAGCCCAGAAGA<br>TGTGGACGCGCTTGAACTCGATCCAGCGATTTCTTATATTGAAGAG<br>GATGCAGAAGTAACGACAATGGCGCAATCAGTGCCATGGGGAATTA<br>GCCGTGTGCAAGCCCCAGCTGCCCATAACCGTGGATTGACAGGTT<br>CTGGTGTAAAAGTTGCTGTCCTCGATACAGGTATTTCCGTTCATCCA<br>GACTTAAATATTCGTGGTGGCGCTAGCTTTGTACCAGGGGAACCAT<br>CCTATCAAGATGGGAATGGGCATGGCACGCATGTGGCCGGGACGA<br>TTGCTGCTTTAGATAATTCGATTGGCGTTCTTGGCGTAGCGCCGAG<br>CGCGGAACTATACGCTGTTAAAGTATTAGGGGCGAGCGGTTCAGGT<br>TCGGTCAGCTCGATTGCCCAAGGATTGGAATGGGCACTTAACAATG<br>GCATGCACGTTGCTAATTTGAGTTTAGGAAGCCCTTCGCCAAGTGC<br>CACACTTGAGCAAGCTGTTAATAGCGCGACTTCTAGAGGCGTTCTT<br>GTTGTAGCGGCATCTGGGAATTCAGGTGCAGGCTCAATCAGCTATC<br>CGGCCCGTTATGCGAACGTTATGGCAGTCGGAGCTGTTGACCAAA<br>ACAACAACCGCGCCAGCTTTTCACAGTATGGCGCAGGGCTTGACA<br>TTGTCGCACCAGGTGTAAACGTGCAGAGCACATACCCAGGTTCAA<br>CGTATGCCAGCTTAAACGGTACATCGATGGCTACTCCTCATGTTGCA<br>GGTGCAGCAGCCCTTGTTAAACAAAAGTATCCATCTTGGTCCAATG<br>TACAAATCCGCAATCATCTAAAGAATACGGCAACGAGCTTAGGAGAT<br>ACGTGGTTGTATGGAAGCGGACTTGTCAATGCAGAAGCGGCAACA<br>CGCCATCACCATCACCATCACCATCACTAA |
| pP43NMK-Sec-aprE2 | TCGCGCGTTTCGGTGATGACGGTGAAAACCTCTGACACATGCAGC<br>TCCCGGAGACGGTCACAGCTTGTCTGTAAGCGGATGCCGGGAGCA<br>GACAAGCCCGTCAGGGCGCGTCAGCGGGTGTTGGCGGGTGTCG<br>GGGCTGGCTTAACTATGCGGCATCAGAGCAGATTGTACTGAGAGTG<br>CACCATATGCGGTGTGAAATACCGCACAGATGCGTAAGGAGAAAAT<br>ACCGCATCAGGCGCCATTCGCCATTCAGGCTGCGCAACTGTTGGG<br>AAGGGCGATCGGTGCGGGCCTCTTCGCTATTACGCCAGCTGGCGA<br>AAGGGGGATGTGCTGCAAGGCGATTAAGTTGGGTAACGCCAGGGT<br>TTTCCCAGTCACGACGTTGTAAAACGACGGCCAGTGAATTCCTTAA<br>GGAACGTACAGACGGCTTAAAAGCCTTTAAAAACGTTTTTAAGGGG<br>TTTGTAGACAAGGTAAAGGATAAAACAGCACAATTCCAAGAAAAACA |

CGATTTAGAACCTAAAAAGAACGAATTTGAACTAACTCATAACCGAG
AGGTAAAAAAAGAACGAAGTCGAGATCAGGGAATGAGTTTATAAAAT
AAAAAAAGCACCTGAAAAGGTGTCTTTTTTTGATGGTTTTGAACTTG
TTCTTTCTTATCTTGATACATATAGAAATAACGTCATTTTTATTTTAGTT
GCTGAAAGGTGCGTTGAAGTGTTGGTATGTATGTGTTTTAAAGTATT
GAAAACCCTTAAAATTGGTTGCACAGAAAAACCCCATCTGTTAAAGT
TATAAGTGACTAAACAAATAACTAAATAGATGGGGGTTTCTTTTAATAT
TATGTGTCCTAATAGTAGCATTTATTCAGATGAAAAATCAAGGGTTTT
AGTGGACAAGACAAAAAGTGGAAAAGTGAGACCATGGAGAGAAAA
GAAAATCGCTAATGTTGATTACTTTGAACTTCTGCATATTCTTGAATT
TAAAAAGGCTGAAAGAGTAAAAGATTGTGCTGAAATATTAGAGTATA
AACAAAATCGTGAAACAGGCGAAAGAAAGTTGTATCGAGTGTGGTT
TTGTAAATCCAGGCTTTGTCCAATGTGCAACTGGAGGAGAGCAATG
AAACATGGCATTCAGTCACAAAAGGTTGTTGCTGAAGTTATTAAACA
AAAGCCAACAGTTCGTTGGTTGTTTCTCACATTAACAGTTAAAAATG
TTTATGATGGCGAAGAATTAAATAAGAGTTTGTCAGATATGGCTCAA
GGATTTCGCCGAATGATGCAATATAAAAAAATTAATAAAAATCTTGTT
GGTTTTATGCGTGCAACGGAAGTGACAATAAATAATAAAGATAATTCT
TATAATCAGCACATGCATGTATTGGTATGTGTGGAACCAACTTATTTT
AAGAATACAGAAAACTACGTGAATCAAAAACAATGGATTCAATTTTG
GAAAAAGGCAATGAAATTAGACTATGATCCAAATGTAAAAGTTCAAA
TGATTCGACCGAAAAATAAATATAAATCGGATATACAATCGGCAATTG
ACGAAACTGCAAAATATCCTGTAAAGGATACGGATTTTATGACCGAT
GATGAAGAAAAGAATTTGAAACGTTTGTCTGATTTGGAGGAAGGTT
TACACCGTAAAAGGTTAATCTCCTATGGTGGTTTGTTAAAAGAAATA
CATAAAAAATTAAACCTTGATGACACAGAAGAAGGCGATTTGATTCA
TACAGATGATGACGAAAAAGCCGATGAAGATGGATTTTCTATTATTG
CAATGTGGAATTGGGAACGGAAAAATTATTTTATTAAAGAGTAGTTC
AACAAACGGGCCAGTTTGTTGAAGATTAGATGCTATAATTGTTATTAA
AAGGATTGAAGGATGCTTAGGAAGACGAGTTATTAATAGCTGAATAA
GAACGGTGCTCTCCAAATATTCTTATTTAGAAAAGCAAATCTAAAATT
ATCTGAAAAGGGAATGAGAATAGTGAATGGACCAATAATAATGACTA
GAGAAGAAAGAATGAAGATTGTTCATGAAATTAAGGAACGAATATTG
GATAAATATGGGGATGATGTTAAGGCTATTGGTGTTTATGGCTCTCTT
GGTCGTCAGACTGATGGGCCCTATTCGGATATTGAGATGATGTGTG
TCATGTCAACAGAGGAAGCAGAGTTCAGCCATGAATGGACAACCG
GTGAGTGGAAGGTGGAAGTGAATTTTGATAGCGAAGAGATTCTACT
AGATTATGCATCTCAGGTGGAATCAGATTGGCCGCTTACACATGGT
CAATTTTTCTCTATTTTGCCGATTTATGATTCAGGTGGATACTTAGAG
AAAGTGTATCAAACTGCTAAATCGGTAGAAGCCCAAACGTTCCACG
ATGCGATTTGTGCCCTTATCGTAGAAGAGCTGTTTGAATATGCAGGC
AAATGGCGTAATATTCGTGTGCAAGGACCGACAACATTTCTACCATC
CTTGACTGTACAGGTAGCAATGGCAGGTGCCATGTTGATTGGTCTG
CATCATCGCATCTGTTATACGACGAGCGCTTCGGTCTTAACTGAAG

CAGTTAAGCAATCAGATCTTCCTTCAGGTTATGACCATCTGTGCCAG
TTCGTAATGTCTGGTCAACTTTCCGACTCTGAGAAACTTCTGGAATC
GCTAGAGAATTTCTGGAATGGGATTCAGGAGTGGACAGAACGACA
CGGATATATAGTGGATGTGTCAAAACGCATACCATTTTGAACGATGA
CCTCTAATAATTGTTAATCATGTTGGTTACGTATTTATTAACTTCTCCT
AGTATTAGTAATTATCATGGCTGTCATGGCGCATTAACGGAATAAAG
GGTGTGCTTAAATCGGGCCATTTTGCGTAATAAGAAAAAGGATTAAT
TATGAGCGAATTGAATTAATAATAAGGTAATAGATTTACATTAGAAAAT
GAAAGGGGATTTTATGCGTGAGAATGTTACAGTCTATCCCGGCATT
GCCAGTCGGGGATATTAAAAAGAGTATAGGTTTTTATTGGGATAAAG
TAGGTTTCACTTTGGTTCACCATGAAGATGGATTCGCAGTTCTAATG
TGTAATGAGGTTCGGATTCATCTATGGGAGGCAAGTGATGAAGGCT
GGCGCCTCGTAGTAATGATTCACCGGTTTGTACAGGTGCGGAGTC
GTTTATTGCTGGTACTGCTAGTTGCCGCATTGAAGTAGAGGGAATT
GATGAATTATATCAACATATTAAGCCTTTGGGCATTTTGCACCCCAAT
ACATCATTAAAAGATCAGTGGTGGGATGAACGAGACTTTGCAGTAAT
TGATCCCGACAACAATTTGATTAGCTTTTTTCAACAAATAAAAAGCTA
AAATCTATTATTAATCTGTTCAGCAATCGGGCGCGATTGCTGAATAAA
AGATACGAGAGACCTCTCTTGTATCTTTTTTATTTTGAGTGGTTTTGT
CCGTTACACTAGAAAACCGAAAGACAATAAAAATTTTATTCTTGCTG
AGTCTGGCTTTCGGTAAGCTAGACAAAACGGACAAAATAAAAATTG
GCAAGGGTTTAAAGGTGGAGATTTTTTGAGTGATCTTCTCAAAAAAT
ACTACCTGTCCCTTGCTGATTTTTAAACGAGCACGAGAGCAAAACC
CCCCTTTGCTGAGGTGGCAGAGGGCAGGTTTTTTTGTTTCTTTTTT
CTCGTAAAAAAAAGAAAGGTCTTAAAGGTTTTATGGTTTTGGTCGGC
ACTGCCGCGCCTCGCAGAGCACACACTTTATGAATATAAAGTATAGT
GTGTTATACTTTACTTGGAAGTGGTTGCCGGAAAGAGCGAAAATGC
CTCACATTTGTGCCACCTAAAAAGGAGCGATTTACATATGAGTTATG
CAGTTTGTAGAATGCAAAAAGTGAAATCAGCTGGACTAAAAGGCAT
GCAATTTCATAATCAAAGAGAGCGAAAAAGTAGAACGAATGATGATA
TTGACCATGAGCGAACACGTGAAAATTATGATTTGAAAAATGATAAA
AATATTGATTACAACGAACGTGTCAAAGAAATTATTGAATCACAAAAA
ACAGGTACAAGAAAAACGAGGAAAGATGCTGTTCTTGTAAATGAGT
TGCTAGTAACATCTGACCGAGATTTTTTTGAGCAACTGGATCCTGAT
AGGTGGTATGTTTTCGCTTGAACTTTTAAATACAGCCATTGAACATA
CGGTTGATTTAATAACTGACAAACATCACCCTCTTGCTAAAGCGGCC
AAGGACGCTGCCGCCGGGGCTGTTTGCGTTTTTGCCGTGATTTCG
TGTATCATTGGTTTACTTATTTTTTTGCCAAAGCTGTAATGGCTGAAA
ATTCTTACATTTATTTTACATTTTTAGAAATGGGCGTGAAAAAAAGCG
CGCGATTATGTAAAATATAAAGTGATAGCGGTACCATTATAGGAGAAG
GAGGAATGTACACATGAAGAAACCGTTGGGGAAAATTGTCGCAAGC
ACCGCACTACTCATTTCTGTTGCTTTTAGTTCATCGATCGCATCGGC
TGCTGAAGAAGCAAAAGAAAAATATTTAATTGGCTTTAATGAGCAGG
AAGCTGTCAGTGAGTTTGTAGAACAAGTAGAGGCAAATGACGAGGT

CGCCATTCTCTCTGAGGAAGAGGAAGTCGAAATTGAATTGCTTCAT
GAATTTGAAACGATTCCTGTTTTATCCGTTGAGTTAAGCCCAGAAGA
TGTGGACGCGCTTGAACTCGATCCAGCGATTTCTTATATTGAAGAG
GATGCAGAAGTAACGACAATGGCGCAATCAGTGCCATGGGGAATTA
GCCGTGTGCAAGCCCCAGCTGCCCATAACCGTGGATTGACAGGTT
CTGGTGTAAAAGTTGCTGTCCTCGATACAGGTATTTCCGTTCATCCA
GACTTAAATATTCGTGGTGGCGCTAGCTTTGTACCAGGGGAACCAT
CCTATCAAGATGGGAATGGGCATGGCACGCATGTGGCCGGGACGA
TTGCTGCTTTAGATAATTCGATTGGCGTTCTTGGCGTAGCGCCGAG
CGCGGAACTATACGCTGTTAAAGTATTAGGGGCGAGCGGTTCAGGT
TCGGTCAGCTCGATTGCCCAAGGATTGGAATGGGCACTTAACAATG
GCATGCACGTTGCTAATTTGAGTTTAGGAAGCCCTTCGCCAAGTGC
CACACTTGAGCAAGCTGTTAATAGCGCGACTTCTAGAGGCGTTCTT
GTTGTAGCGGCATCTGGGAATTCAGGTGCAGGCTCAATCAGCTATC
CGGCCCGTTATGCGAACGTTATGGCAGTCGGAGCTGTTGACCAAA
ACAACAACCGCGCCAGCTTTTCACAGTATGGCGCAGGGCTTGACA
TTGTCGCACCAGGTGTAAACGTGCAGAGCACATACCCAGGTTCAA
CGTATGCCAGCTTAAACGGTACATCGATGGCTACTCCTCATGTTGCA
GGTGCAGCAGCCCTTGTTAAACAAAAGTATCCATCTTGGTCCAATG
TACAAATCCGCAATCATCTAAAGAATACGGCAACGAGCTTAGGAGAT
ACGTGGTTGTATGGAAGCGGACTTGTCAATGCAGAAGCGGCAACA
CGCCATCACCATCACCATCACCATCACTAATGATGAAAGCTTGGCG
TAATCATGGTCATAGCTGTTTCCTGTGTGAAATTGTTATCCGCTCAC
AATTCCACACAACATACGAGCCGGAAGCATAAAGTGTAAAGCCTGG
GGTGCCTAATGAGTGAGCTAACTCACATTAATTGCGTTGCGCTCAC
TGCCCGCTTTCCAGTCGGGAAACCTGTCGTGCCAGCTGCATTAAT
GAATCGGCCAACGCGCGGGGAGAGGCGGTTTGCGTATTGGGCGC
TCTTCCGCTTCCTCGCTCACTGACTCGCTGCGCTCGGTCGTTCGG
CTGCGGCGAGCGGTATCAGCTCACTCAAAGGCGGTAATACGGTTAT
CCACAGAATCAGGGGATAACGCAGGAAAGAACATGTGAGCAAAAG
GCCAGCAAAAGGCCAGGAACCGTAAAAAGGCCGCGTTGCTGGCG
TTTTTCCATAGGCTCCGCCCCCCTGACGAGCATCACAAAAATCGAC
GCTCAAGTCAGAGGTGGCGAAACCCGACAGGACTATAAAGATACC
AGGCGTTTCCCCCTGGAAGCTCCCTCGTGCGCTCTCCTGTTCCGA
CCCTGCCGCTTACCGGATACCTGTCCGCCTTTCTCCCTTCGGGAA
GCGTGGCGCTTTCTCATAGCTCACGCTGTAGGTATCTCAGTTCGGT
GTAGGTCGTTCGCTCCAAGCTGGGCTGTGTGCACGAACCCCCCGT
TCAGCCCGACCGCTGCGCCTTATCCGGTAACTATCGTCTTGAGTCC
AACCCGGTAAGACACGACTTATCGCCACTGGCAGCAGCCACTGGT
AACAGGATTAGCAGAGCGAGGTATGTAGGCGGTGCTACAGAGTTCT
TGAAGTGGTGGCCTAACTACGGCTACACTAGAAGAACAGTATTTGG
TATCTGCGCTCTGCTGAAGCCAGTTACCTTCGGAAAAAGAGTTGGT
AGCTCTTGATCCGGCAAACAAACCACCGCTGGTAGCGGTGGTTTTT
TTGTTTGCAAGCAGCAGATTACGCGCAGAAAAAAAGGATCTCAAGA

| | |
|---|---|
| | AGATCCTTTGATCTTTTCTACGGGGTCTGACGCTCAGTGGAACGAA<br>AACTCACGTTAAGGGATTTTGGTCATGAGATTATCAAAAAGGATCTT<br>CACCTAGATCCTTTTAAATTAAAAATGAAGTTTTAAATCAATCTAAAGT<br>ATATATGAGTAAACTTGGTCTGACAGTTACCAATGCTTAATCAGTGA<br>GGCACCTATCTCAGCGATCTGTCTATTTCGTTCATCCATAGTTGCCT<br>GACTCCCCGTCGTGTAGATAACTACGATACGGGAGGGCTTACCATC<br>TGGCCCCAGTGCTGCAATGATACCGCGAGACCCACGCTCACCGGC<br>TCCAGATTTATCAGCAATAAACCAGCCAGCCGGAAGGGCCGAGCG<br>CAGAAGTGGTCCTGCAACTTTATCCGCCTCCATCCAGTCTATTAATT<br>GTTGCCGGGAAGCTAGAGTAAGTAGTTCGCCAGTTAATAGTTTGCG<br>CAACGTTGTTGCCATTGCTACAGGCATCGTGGTGTCACGCTCGTC<br>GTTTGGTATGGCTTCATTCAGCTCCGGTTCCCAACGATCAAGGCGA<br>GTTACATGATCCCCCATGTTGTGCAAAAAAGCGGTTAGCTCCTTCG<br>GTCCTCCGATCGTTGTCAGAAGTAAGTTGGCCGCAGTGTTATCACT<br>CATGGTTATGGCAGCACTGCATAATTCTCTTACTGTCATGCCATCCG<br>TAAGATGCTTTTCTGTGACTGGTGAGTACTCAACCAAGTCATTCTGA<br>GAATAGTGTATGCGGCGACCGAGTTGCTCTTGCCCGGCGTCAATAC<br>GGGATAATACCGCGCCACATAGCAGAACTTTAAAAGTGCTCATCATT<br>GGAAAACGTTCTTCGGGGCGAAAACTCTCAAGGATCTTACCGCTGT<br>TGAGATCCAGTTCGATGTAACCCACTCGTGCACCCAACTGATCTTC<br>AGCATCTTTTACTTTCACCAGCGTTTCTGGGTGAGCAAAAACAGGA<br>AGGCAAAATGCCGCAAAAAAGGGAATAAGGGCGACACGGAAATGT<br>TGAATACTCATACTCTTCCTTTTTCAATATTATTGAAGCATTTATCAGG<br>GTTATTGTCTCATGAGCGGATACATATTTGAATGTATTTAGAAAAATAA<br>ACAAATAGGGGTTCCGCGCACATTTCCCCGAAAAGTGCCACCTGA<br>CGTCTAAGAAACCATTATTATCATGACATTAACCTATAAAAATAGGCG<br>TATCACGAGGCCCTTTCGTC |
| pP43NMK-Tat-aprE2 | TCGCGCGTTTCGGTGATGACGGTGAAAACCTCTGACACATGCAGC<br>TCCCGGAGACGGTCACAGCTTGTCTGTAAGCGGATGCCGGGAGCA<br>GACAAGCCCGTCAGGGCGCGTCAGCGGGTGTTGGCGGGTGTCG<br>GGGCTGGCTTAACTATGCGGCATCAGAGCAGATTGTACTGAGAGTG<br>CACCATATGCGGTGTGAAATACCGCACAGATGCGTAAGGAGAAAAT<br>ACCGCATCAGGCGCCATTCGCCATTCAGGCTGCGCAACTGTTGGG<br>AAGGGCGATCGGTGCGGGCCTCTTCGCTATTACGCCAGCTGGCGA<br>AAGGGGGATGTGCTGCAAGGCGATTAAGTTGGGTAACGCCAGGGT<br>TTTCCCAGTCACGACGTTGTAAAACGACGGCCAGTGAATTCCTTAA<br>GGAACGTACAGACGGCTTAAAAGCCTTTAAAAACGTTTTTAAGGGG<br>TTTGTAGACAAGGTAAAGGATAAAACAGCACAATTCCAAGAAAAACA<br>CGATTTAGAACCTAAAAAGAACGAATTTGAACTAACTCATAACCGAG<br>AGGTAAAAAAAGAACGAAGTCGAGATCAGGGAATGAGTTTATAAAAT<br>AAAAAAAGCACCTGAAAAGGTGTCTTTTTTTGATGGTTTTGAACTTG<br>TTCTTTCTTATCTTGATACATATAGAAATAACGTCATTTTTATTTTAGTT<br>GCTGAAAGGTGCGTTGAAGTGTTGGTATGTATGTGTTTTAAAGTATT<br>GAAAACCCTTAAAATTGGTTGCACAGAAAAACCCCATCTGTTAAAGT |

TATAAGTGACTAAACAAATAACTAAATAGATGGGGGTTTCTTTTAATAT
TATGTGTCCTAATAGTAGCATTTATTCAGATGAAAAATCAAGGGTTTT
AGTGGACAAGACAAAAAGTGGAAAAGTGAGACCATGGAGAGAAAA
GAAAATCGCTAATGTTGATTACTTTGAACTTCTGCATATTCTTGAATT
TAAAAAGGCTGAAAGAGTAAAAGATTGTGCTGAAATATTAGAGTATA
AACAAAATCGTGAAACAGGCGAAAGAAAGTTGTATCGAGTGTGGTT
TTGTAAATCCAGGCTTTGTCCAATGTGCAACTGGAGGAGAGCAATG
AAACATGGCATTCAGTCACAAAAGGTTGTTGCTGAAGTTATTAAACA
AAAGCCAACAGTTCGTTGGTTGTTTCTCACATTAACAGTTAAAAATG
TTTATGATGGCGAAGAATTAAATAAGAGTTTGTCAGATATGGCTCAA
GGATTTCGCCGAATGATGCAATATAAAAAAATTAATAAAAATCTTGTT
GGTTTTATGCGTGCAACGGAAGTGACAATAAATAATAAAGATAATTCT
TATAATCAGCACATGCATGTATTGGTATGTGTGGAACCAACTTATTTT
AAGAATACAGAAAACTACGTGAATCAAAAACAATGGATTCAATTTTG
GAAAAAGGCAATGAAATTAGACTATGATCCAAATGTAAAAGTTCAAA
TGATTCGACCGAAAAATAAATATAAATCGGATATACAATCGGCAATTG
ACGAAACTGCAAAATATCCTGTAAAGGATACGGATTTTATGACCGAT
GATGAAGAAAAGAATTTGAAACGTTTGTCTGATTTGGAGGAAGGTT
TACACCGTAAAAGGTTAATCTCCTATGGTGGTTTGTTAAAAGAAATA
CATAAAAAATTAAACCTTGATGACACAGAAGAAGGCGATTTGATTCA
TACAGATGATGACGAAAAAGCCGATGAAGATGGATTTTCTATTATTG
CAATGTGGAATTGGGAACGGAAAAATTATTTTATTAAAGAGTAGTTC
AACAAACGGGCCAGTTTGTTGAAGATTAGATGCTATAATTGTTATTAA
AAGGATTGAAGGATGCTTAGGAAGACGAGTTATTAATAGCTGAATAA
GAACGGTGCTCTCCAAATATTCTTATTTAGAAAAGCAAATCTAAAATT
ATCTGAAAAGGGAATGAGAATAGTGAATGGACCAATAATAATGACTA
GAGAAGAAAGAATGAAGATTGTTCATGAAATTAAGGAACGAATATTG
GATAAATATGGGGATGATGTTAAGGCTATTGGTGTTTATGGCTCTCTT
GGTCGTCAGACTGATGGGCCCTATTCGGATATTGAGATGATGTGTG
TCATGTCAACAGAGGAAGCAGAGTTCAGCCATGAATGGACAACCG
GTGAGTGGAAGGTGGAAGTGAATTTTGATAGCGAAGAGATTCTACT
AGATTATGCATCTCAGGTGGAATCAGATTGGCCGCTTACACATGGT
CAATTTTTCTCTATTTTGCCGATTTATGATTCAGGTGGATACTTAGAG
AAAGTGTATCAAACTGCTAAATCGGTAGAAGCCCAAACGTTCCACG
ATGCGATTTGTGCCCTTATCGTAGAAGAGCTGTTTGAATATGCAGGC
AAATGGCGTAATATTCGTGTGCAAGGACCGACAACATTTCTACCATC
CTTGACTGTACAGGTAGCAATGGCAGGTGCCATGTTGATTGGTCTG
CATCATCGCATCTGTTATACGACGAGCGCTTCGGTCTTAACTGAAG
CAGTTAAGCAATCAGATCTTCCTTCAGGTTATGACCATCTGTGCCAG
TTCGTAATGTCTGGTCAACTTTCCGACTCTGAGAAACTTCTGGAATC
GCTAGAGAATTTCTGGAATGGGATTCAGGAGTGGACAGAACGACA
CGGATATATAGTGGATGTGTCAAAACGCATACCATTTTGAACGATGA
CCTCTAATAATTGTTAATCATGTTGGTTACGTATTTATTAACTTCTCCT
AGTATTAGTAATTATCATGGCTGTCATGGCGCATTAACGGAATAAAG

GGTGTGCTTAAATCGGGCCATTTTGCGTAATAAGAAAAAGGATTAAT
TATGAGCGAATTGAATTAATAATAAGGTAATAGATTTACATTAGAAAAT
GAAAGGGGATTTTATGCGTGAGAATGTTACAGTCTATCCCGGCATT
GCCAGTCGGGGATATTAAAAAGAGTATAGGTTTTTATTGGGATAAAG
TAGGTTTCACTTTGGTTCACCATGAAGATGGATTCGCAGTTCTAATG
TGTAATGAGGTTCGGATTCATCTATGGGAGGCAAGTGATGAAGGCT
GGCGCCTCGTAGTAATGATTCACCGGTTTGTACAGGTGCGGAGTC
GTTTATTGCTGGTACTGCTAGTTGCCGCATTGAAGTAGAGGGAATT
GATGAATTATATCAACATATTAAGCCTTTGGGCATTTTGCACCCCAAT
ACATCATTAAAAGATCAGTGGTGGGATGAACGAGACTTTGCAGTAAT
TGATCCCGACAACAATTTGATTAGCTTTTTTCAACAAATAAAAAGCTA
AAATCTATTATTAATCTGTTCAGCAATCGGGCGCGATTGCTGAATAAA
AGATACGAGAGACCTCTCTTGTATCTTTTTTATTTTGAGTGGTTTTGT
CCGTTACACTAGAAAACCGAAAGACAATAAAAATTTTATTCTTGCTG
AGTCTGGCTTTCGGTAAGCTAGACAAAACGGACAAAATAAAAATTG
GCAAGGGTTTAAAGGTGGAGATTTTTTGAGTGATCTTCTCAAAAAAT
ACTACCTGTCCCTTGCTGATTTTTAAACGAGCACGAGAGCAAAACC
CCCCTTTGCTGAGGTGGCAGAGGGCAGGTTTTTTTGTTTCTTTTTT
CTCGTAAAAAAAAGAAAGGTCTTAAAGGTTTTATGGTTTTGGTCGGC
ACTGCCGCGCCTCGCAGAGCACACACTTTATGAATATAAAGTATAGT
GTGTTATACTTTACTTGGAAGTGGTTGCCGGAAAGAGCGAAAATGC
CTCACATTTGTGCCACCTAAAAAGGAGCGATTTACATATGAGTTATG
CAGTTTGTAGAATGCAAAAAGTGAAATCAGCTGGACTAAAAGGCAT
GCAATTTCATAATCAAAGAGAGCGAAAAAGTAGAACGAATGATGATA
TTGACCATGAGCGAACACGTGAAAATTATGATTTGAAAAATGATAAA
AATATTGATTACAACGAACGTGTCAAAGAAATTATTGAATCACAAAAA
ACAGGTACAAGAAAAACGAGGAAAGATGCTGTTCTTGTAAATGAGT
TGCTAGTAACATCTGACCGAGATTTTTTTGAGCAACTGGATCCTGAT
AGGTGGTATGTTTTCGCTTGAACTTTTAAATACAGCCATTGAACATA
CGGTTGATTTAATAACTGACAAACATCACCCTCTTGCTAAAGCGGCC
AAGGACGCTGCCGCCGGGGCTGTTTGCGTTTTTGCCGTGATTTCG
TGTATCATTGGTTTACTTATTTTTTTGCCAAAGCTGTAATGGCTGAAA
ATTCTTACATTTATTTTACATTTTTAGAAATGGGCGTGAAAAAAAGCG
CGCGATTATGTAAAATATAAAGTGATAGCGGTACCATTATAGGAGAAG
GAGGAATGTACACATGGCATATGATAGTCGCTTTGACGAGTGGGTG
CAAAAGCTTAAGGAAGAGTCATTCCAGAACAATACGTTCGATCGGC
GTAAGTTTATCCAGGGTGCAGGAAAAATTGCAGGACTTAGCCTGGG
TCTGACAATTGCCCAAAGTGTGGGCGCCTTTGAAGTGAACGCCGC
TGAAGAAGCAAAAGAAAAATATTTAATTGGCTTTAATGAGCAGGAAG
CTGTCAGTGAGTTTGTAGAACAAGTAGAGGCAAATGACGAGGTCG
CCATTCTCTCTGAGGAAGAGGAAGTCGAAATTGAATTGCTTCATGA
ATTTGAAACGATTCCTGTTTTATCCGTTGAGTTAAGCCCAGAAGATG
TGGACGCGCTTGAACTCGATCCAGCGATTTCTTATATTGAAGAGGAT
GCAGAAGTAACGACAATGGCGCAATCAGTGCCATGGGGAATTAGC

CGTGTGCAAGCCCCAGCTGCCCATAACCGTGGATTGACAGGTTCT
GGTGTAAAAGTTGCTGTCCTCGATACAGGTATTTCCGTTCATCCAGA
CTTAAATATTCGTGGTGGCGCTAGCTTTGTACCAGGGGAACCATCC
TATCAAGATGGGAATGGGCATGGCACGCATGTGGCCGGGACGATT
GCTGCTTTAGATAATTCGATTGGCGTTCTTGGCGTAGCGCCGAGCG
CGGAACTATACGCTGTTAAAGTATTAGGGGCGAGCGGTTCAGGTTC
GGTCAGCTCGATTGCCCAAGGATTGGAATGGGCACTTAACAATGGC
ATGCACGTTGCTAATTTGAGTTTAGGAAGCCCTTCGCCAAGTGCCA
CACTTGAGCAAGCTGTTAATAGCGCGACTTCTAGAGGCGTTCTTGT
TGTAGCGGCATCTGGGAATTCAGGTGCAGGCTCAATCAGCTATCCG
GCCCGTTATGCGAACGTTATGGCAGTCGGAGCTGTTGACCAAAACA
ACAACCGCGCCAGCTTTTCACAGTATGGCGCAGGGCTTGACATTG
TCGCACCAGGTGTAAACGTGCAGAGCACATACCCAGGTTCAACGTA
TGCCAGCTTAAACGGTACATCGATGGCTACTCCTCATGTTGCAGGT
GCAGCAGCCCTTGTTAAACAAAAGTATCCATCTTGGTCCAATGTACA
AATCCGCAATCATCTAAAGAATACGGCAACGAGCTTAGGAGATACGT
GGTTGTATGGAAGCGGACTTGTCAATGCAGAAGCGGCAACACGCC
ATCACCATCACCATCACCATCACTAATGATGAAAGCTTGGCGTAATC
ATGGTCATAGCTGTTTCCTGTGTGAAATTGTTATCCGCTCACAATTC
CACACAACATACGAGCCGGAAGCATAAAGTGTAAAGCCTGGGGTG
CCTAATGAGTGAGCTAACTCACATTAATTGCGTTGCGCTCACTGCC
CGCTTTCCAGTCGGGAAACCTGTCGTGCCAGCTGCATTAATGAATC
GGCCAACGCGCGGGGAGAGGCGGTTTGCGTATTGGGCGCTCTTC
CGCTTCCTCGCTCACTGACTCGCTGCGCTCGGTCGTTCGGCTGCG
GCGAGCGGTATCAGCTCACTCAAAGGCGGTAATACGGTTATCCACA
GAATCAGGGGATAACGCAGGAAAGAACATGTGAGCAAAAGGCCAG
CAAAAGGCCAGGAACCGTAAAAAGGCCGCGTTGCTGGCGTTTTTC
CATAGGCTCCGCCCCCCTGACGAGCATCACAAAAATCGACGCTCA
AGTCAGAGGTGGCGAAACCCGACAGGACTATAAAGATACCAGGCG
TTTCCCCCTGGAAGCTCCCTCGTGCGCTCTCCTGTTCCGACCCTG
CCGCTTACCGGATACCTGTCCGCCTTTCTCCCTTCGGGAAGCGTG
GCGCTTTCTCATAGCTCACGCTGTAGGTATCTCAGTTCGGTGTAGG
TCGTTCGCTCCAAGCTGGGCTGTGTGCACGAACCCCCCGTTCAGC
CCGACCGCTGCGCCTTATCCGGTAACTATCGTCTTGAGTCCAACCC
GGTAAGACACGACTTATCGCCACTGGCAGCAGCCACTGGTAACAG
GATTAGCAGAGCGAGGTATGTAGGCGGTGCTACAGAGTTCTTGAAG
TGGTGGCCTAACTACGGCTACACTAGAAGAACAGTATTTGGTATCTG
CGCTCTGCTGAAGCCAGTTACCTTCGGAAAAAGAGTTGGTAGCTCT
TGATCCGGCAAACAAACCACCGCTGGTAGCGGTGGTTTTTTTGTTT
GCAAGCAGCAGATTACGCGCAGAAAAAAAGGATCTCAAGAAGATCC
TTTGATCTTTTCTACGGGGTCTGACGCTCAGTGGAACGAAAACTCA
CGTTAAGGGATTTTGGTCATGAGATTATCAAAAAGGATCTTCACCTA
GATCCTTTTAAATTAAAAATGAAGTTTTAAATCAATCTAAAGTATATAT
GAGTAAACTTGGTCTGACAGTTACCAATGCTTAATCAGTGAGGCAC

CTATCTCAGCGATCTGTCTATTTCGTTCATCCATAGTTGCCTGACTC
CCCGTCGTGTAGATAACTACGATACGGGAGGGCTTACCATCTGGCC
CCAGTGCTGCAATGATACCGCGAGACCCACGCTCACCGGCTCCAG
ATTTATCAGCAATAAACCAGCCAGCCGGAAGGGCCGAGCGCAGAA
GTGGTCCTGCAACTTTATCCGCCTCCATCCAGTCTATTAATTGTTGC
CGGGAAGCTAGAGTAAGTAGTTCGCCAGTTAATAGTTTGCGCAACG
TTGTTGCCATTGCTACAGGCATCGTGGTGTCACGCTCGTCGTTTGG
TATGGCTTCATTCAGCTCCGGTTCCCAACGATCAAGGCGAGTTACA
TGATCCCCCATGTTGTGCAAAAAAGCGGTTAGCTCCTTCGGTCCTC
CGATCGTTGTCAGAAGTAAGTTGGCCGCAGTGTTATCACTCATGGT
TATGGCAGCACTGCATAATTCTCTTACTGTCATGCCATCCGTAAGAT
GCTTTTCTGTGACTGGTGAGTACTCAACCAAGTCATTCTGAGAATA
GTGTATGCGGCGACCGAGTTGCTCTTGCCCGGCGTCAATACGGGA
TAATACCGCGCCACATAGCAGAACTTTAAAAGTGCTCATCATTGGAA
AACGTTCTTCGGGGCGAAAACTCTCAAGGATCTTACCGCTGTTGAG
ATCCAGTTCGATGTAACCCACTCGTGCACCCAACTGATCTTCAGCA
TCTTTTACTTTCACCAGCGTTTCTGGGTGAGCAAAAACAGGAAGGC
AAAATGCCGCAAAAAAGGGAATAAGGGCGACACGGAAATGTTGAAT
ACTCATACTCTTCCTTTTTCAATATTATTGAAGCATTTATCAGGGTTAT
TGTCTCATGAGCGGATACATATTTGAATGTATTTAGAAAAATAAACAA
ATAGGGGTTCCGCGCACATTTCCCCGAAAAGTGCCACCTGACGTC
TAAGAAACCATTATTATCATGACATTAACCTATAAAAATAGGCGTATCA
CGAGGCCCTTTCGTC

**Table S3.** Signal peptides protein sequences in optimization of expression experiment

| Names | Sequence |
|---|---|
| **ApGA signal peptides** | |
| α-factor | MRFPSIFTAVLFAASSALA |
| AE1 | MKLFFTLLVFASTVFG |
| AE2 | MKSFFLLALLSATVFA |
| AE3 | MRGLSLLAVLTLLAATAFA |
| AE4 | MKLFSFLTILFLSSVFALTTNA |
| AE5 | MKVSLLLATFFSSVLA |
| AE6 | MKLLLFVFAFLASVSA |
| AE7 | MKHLLVLLLAVSLVASA |
| AE8 | MKSNFAVFVVFSLLLVANLSCA |
| AE9 | MKLALLLSLAAPALA |
| AE10 | MKLVLTALLAASVFA |
| **aprE2 signal peptides** | |
| AprE2_SP | MKKPLGKIVASTALLISVAFSSSIASA |
| AS1 | MKKLLSLLLASTVFFSGLAFA |
| AS2 | MKKFLLLSTLFFLSTSVFA |
| AS3 | MKKVLLLSSILFSLFSFSAFA |
| AS4 | MKKILFSLSLLFSFSAFA |
| AS5 | MKKSLLSLFAASLLTSSAFA |
| AS6 | MKKIFLSLVLFFSGLALA |
| AS7 | MRIVAAALLMSSLLFGGLLSAGPAQA |
| AS8 | MKKTLLSGALAAASLLATASPALA |
| AS9 | MKKLLLAALGATLGLAFAAPASA |
| AS10 | MRKRLLLAAALAVAALSAAAPASA |
| PhoD | AGAAGTAGAAGTAGCTCTAGTAGTAGTGG |
| AT1 | MKRREFLKLSVLTAGTAASLSLSALPALA |
| AT2 | MNRREFLKRSAALSLLAASSLPLGASSAQA |
| AT3 | MNRREFLKGLSAGAASGLAGLALSSSAAMA |
| AT4 | MERRTFLKLAGALAAGVGAGALLAGTPAKA |
| AT5 | MTRREFLKSAAASALAAAPLTASA |
| AT6 | MTRREFLLAGAAASGAMAAALLPGAQA |
| AT7 | MSRRKFLKRRSLLGAAGLGLTAAASGPALA |
| AT8 | MSRLSRRGFTRREFLTGAGLLSASALASTPALA |
| AT9 | MNRRKFLKTLGSAALGLAATALAPLPALA |
| AT10 | MGRREFLGTAGALATAALATPALA |

**Table S4.** DNA primers used in replacement of signal peptides

| Names | Sequence (5'-3') |
| --- | --- |
| **ApGA signal peptides primers** | |
| F-AE1 | CTGCTTGTTTTCGCATCTACTGTTTTTGGAGCTACTGGTTCTCTGTCTAGTTGGTTG |
| R-AE1 | GTAGATGCGAAAACAAGCAGGGTGAAGAATAACTTAGCTAATGCGGAGGATGCTGC |
| F-AE2 | GTTGGCATTGCTGAGTGCTACTGTTTTCGCTGCTACTGGTTCTCTGTCTAGTTGGTTG |
| R-AE2 | GTAGCACTCAGCAATGCCAACAGAAAAAAGGACTTAGCTAATGCGGAGGATGCTGC |
| F-AE3 | CTGTTTTGACTCTGTTGGCTGCTACAGCATTCGCTGCTACTGGTTCTCTGTCTAGTTGGTTG |
| R-AE3 | GCCAACAGAGTCAAAACAGCCAAAAGGGACAAACCTCTAGCTAATGCGGAGGATGCTGC |
| F-AE4 | CTTGTTCCTGTCTTCCGTCTTTGCTTTGACCACCAATGCTGCTACTGGTTCTCTGTCTAGTTGGTT |
| R-AE4 | GACGGAAGACAGGAACAAGATTGTCAAGAAGGAGAACAGTTTAGCTAATGCGGAGGATGCTGC |
| F-AE5 | GGCTACTTTCTTCTCTTCCGTTTTGGCTGCTACTGGTTCTCTGTCTAGTTGGTTG |
| R-AE5 | CGGAAGAGAAGAAAGTAGCCAACAACAAACTAACCTTAGCTAATGCGGAGGATGCTGC |
| F-AE6 | CGTGTTCGCATTTCTTGCTAGTGTTTCTGCTGCTACTGGTTCTCTGTCTAGTTGGTTG |
| R-AE6 | CTAGCAAGAAATGCGAACACGAACAGCAACAACTTAGCTAATGCGGAGGATGCTGC |
| F-AE7 | CTTCTTTTGGCTGTTTCATTGGTTGCTTCTGCTGCTACTGGTTCTCTGTCTAGTTGGTTG |
| R-AE7 | CAATGAAACAGCCAAAAGAAGAACCAACAAATGCTTAGCTAATGCGGAGGATGCTGC |
| F-AE8 | GTGTTCTCTTTGCTTTTGGTAGCTAATCTGTCTTGCGCCGCTACTGGTTCTCTGTCTAGTTGGTTG |
| R-AE8 | CTACCAAAAGCAAAGAGAACACGACGAAAACAGCGAAGTTAGACTTAGCTAATGCGGAGGATGCTGC |
| F-AE9 | GCTTAGTCTGGCTGCTCCAGCTTTGGCTGCTACTGGTTCTCTGTCTAGTTGGTTG |
| R-AE9 | CTGGAGCAGCCAGACTAAGCAAAAGAGCAAGTTTAGCTAATGCGGAGGATGCTGC |
| F-AE10 | CTGCTCTGCTGGCCGCTAGTGTTTTCGCTGCTACTGGTTCTCTGTCTAGTTGGTTG |
| R-AE10 | CACTAGCGGCCAGCAGAGCAGTCAAAACCAACTTAGCTAATGCGGAGGATGCTGC |

| | |
|---|---|
| F-ApGA-signalP-VF | CGACTGGTTCCAATTGACAAGCTTTTG |
| R-ApGA-signalP-VF | GTTCCAAACTCTTATTACCGGCGATCAA |
| **aprE2 signal peptides primers** | |
| F-AS1 | GGCCTCTACAGTGTTTTTCAGCGGATTAGCATTTGCAGCTGAAGAAGCAAAAGAAAAATATTTAATTGGCT |
| R-AS1 | AAAAACACTGTAGAGGCCAAAAGTAAGCTCAAGAGTTTCTTCATGTGTACATTCCTCCTTCTCCTATAATGGTACC |
| F-AS2 | GAGCACGCTTTTCTTTCTTAGCACCTCTGTGTTTGCTGCTGAAGAAGCAAAAGAAAAATATTTAATTGGCT |
| R-AS2 | TAAGAAAGAAAAGCGTGCTCAAAAGCAGGAATTTTTTCATGTGTACATTCCTCCTTCTCCTATAATGGTACC |
| F-AS3 | AATCCTGTTTAGCCTGTTTAGCTTTAGTGCGTTCGCCGCTGAAGAAGCAAAAGAAAAATATTTAATTGGCT |
| R-AS3 | AACAGGCTAAACAGGATTGAGCTAAGTAACAGGACTTTCTTCATGTGTACATTCCTCCTTCTCCTATAATGGTACC |
| F-AS4 | CTCCCTGTCTCTTCTGTTCAGCTTTTCCGCTTTTGCCGCTGAAGAAGCAAAAGAAAAATATTTAATTGGCT |
| R-AS4 | CTGAACAGAAGAGACAGGGAGAAGAGGATTTTCTTCATGTGTACATTCCTCCTTCTCCTATAATGGTACC |
| F-AS5 | CTTATTTGCGGCTAGCTTACTGACATCCAGCGCGTTTGCTGCTGAAGAAGCAAAAGAAAAATATTTAATTGGCT |
| R-AS5 | GTAAGCTAGCCGCAAATAAGCTCAGCAGGCTTTTTTTCATGTGTACATTCCTCCTTCTCCTATAATGGTACC |
| F-AS6 | CTTGGTGCTGTTCTTTTCGGGACTCGCACTTGCCGCTGAAGAAGCAAAAGAAAAATATTTAATTGGCT |
| R-AS6 | GAAAAGAACAGCACCAAGGAGAGGAAGATTTTCTTCATGTGTACATTCCTCCTTCTCCTATAATGGTACC |
| F-AS7 | GAGCTCACTTCTTTTTGGCGGATTGTTAAGCGCTGGCCCGGCACAGGCAGCTGAAGAAGCAAAAGAAAAATATTTAATTGGCT |
| R-AS7 | GCCAAAAAGAAGTGAGCTCATCAGCAAGGCGGCCGCAACAATTCTCATGTGTACATTCCTCCTTCTCCTATAATGGTACC |
| F-AS8 | TTGCAGCAGCATCATTGCTTGCAACAGCGAGCCCGGCGCTGGCAGCTGAAGAAGCAAAAGAAAAATATTTAATTGGCT |
| R-AS8 | CAATGATGCTGCTGCAAGTGCTCCACTCAGCAGGGTTTTTTTCATGTGTACATTCCTCCTTCTCCTATAATGGTACC |
| F-AS9 | CTTGGCGCAACGCTGGGCTTAGCATTTGCGGCTCCTGCATCGGCAGCTGAAGAAGCAAAAGAAAAATATTTAATTGGCT |
| R-AS9 | AAGCCCAGCGTTGCGCCAAGAGCTGCAAGAAGCAGTTTCTTCATGTGTACATTCCTCCTTCTCCTATAATGGTACC |
| F-AS10 | GGCGTTGGCGGTGGCTGCGTTGTCAGCAGCAGCCCCGGCATCAGCTGCTGAAGAAGCAAAAGAAAAATATTTAATTGGCT |
| R-AS10 | CGCAGCCACCGCCAACGCCGCCGCCAGCAGTAAGCGTTTTCTC |

| | |
|---|---|
| | ATGTGTACATTCCTCCTTCTCCTATAATGGTA |
| F-AT1 | CAGCAGGAACCGCCGCAAGCCTGTCTCTGAGCGCATTGCCAGCGTTAGCCGCTGAAGAAGCAAAAGAAAAATATTTAATTGGCTTTAA |
| R-AT1 | CTTGCGGCGGTTCCTGCTGTCAAGACGCTCAGTTTCAGGAATTCGCGACGTTTCATGTGTACATTCCTCCTTCTCCTATAATGGTACC |
| F-AT2 | ATCGTTACTTGCAGCATCATCTCTGCCGCTGGGAGCCTCATCAGCGCAGGCCGCTGAAGAAGCAAAAGAAAAATATTTAATTGGCTTTA |
| R-AT2 | ATGATGCTGCAAGTAACGATAATGCCGCTGAGCGCTTCAGAAATTCCCGGCGATTCATGTGTACATTCCTCCTTCTCCTATAATGGTAC |
| F-AT3 | CTGGCGCCGCGTCAGGATTGGCGGGCCTTGCGTTATCATCAAGCGCTGCAATGGCGGCTGAAGAAGCAAAAGAAAAATATTTAATTGGCTTT |
| R-AT3 | CAATCCTGACGCGGCGCCAGCTGACAAGCCTTTAAGGAATTCCCGTCTGTTCATGTGTACATTCCTCCTTCTCCTATAATGGTACC |
| F-AT4 | CCGCAGGAGTTGGAGCGGGAGCCCTTCTTGCTGGAACACCGGCCAAAGCAGCTGAAGAAGCAAAAGAAAAATATTTAATTGGCTTTAA |
| R-AT4 | CCGCTCCAACTCCTGCGGCAAGAGCTCCAGCCAATTTCAGGAAAGTCCGTCTTTCCATGTGTACATTCCTCCTTCTCCTATAATGGTAC |
| F-AT5 | AGCATCAGCATTAGCAGCTGCTCCGCTGACAGCTAGCGCAGCTGAAGAAGCAAAAGAAAAATATTTAATTGGCTTTAATG |
| R-AT5 | CAGCTGCTAATGCTGATGCTGCTGCAGATTTGAGGAATTCTCTGCGTGTCATGTGTACATTCCTCCTTCTCCTATAATGGTAC |
| F-AT6 | CAGCGGCCTCTGGAGCAATGGCTGCGGCCTTGTTGCCGGGAGCACAAGCCGCTGAAGAAGCAAAAGAAAAATATTTAATTGGCTTTAA |
| R-AT6 | CATTGCTCCAGAGGCCGCTGCGCCCGCAAGAAGAAATTCGCGTCTCGTCATGTGTACATTCCTCCTTCTCCTATAATGGTACC |
| F-AT7 | GCGCCGCTGGCTTAGGTCTGACAGCAGCTGCGAGCGGTCCGGCGCTGGCTGCTGAAGAAGCAAAAGAAAAATATTTAATTGGCTTTAA |
| R-AT7 | AGACCTAAGCCAGCGGCGCCTAACAATGAACGGCGCTTCAGAAATTTACGGCGAGACATGTGTACATTCCTCCTTCTCCTATAATGGTA |
| F-AT8 | TACAGGAGCTGGATTGCTGTCAGCGTCAGCACTGGCCAGCACACCGGCACTGGCGGCTGAAGAAGCAAAAGAAAAATATTTAATTGGCT |
| R-AT8 | CAGCAATCCAGCTCCTGTAAGAAATTCGCGGCGCGTAAAGCCTCTTCTAGATAACCGGGACATGTGTACATTCCTCCTTCTCCTATAATGGTACC |
| F-AT9 | AGCACTGGGTTTAGCCGCGACTGCTTTAGCACCGCTGCCAGCATTAGCCGCTGAAGAAGCAAAAGAAAAATATTTAATTGGCTTTAATG |
| R-AT9 | CGCGGCTAAACCCAGTGCTGCGCTTCCCAACGTTTTCAGGAATTTCCGGCGGTTCATGTGTACATTCCTCCTTCTCCTATAATGGTACC |
| F-AT10 | GGCACAGCAGGAGCGCTTGCTACTGCTGCACTTGCAACACCGGCCTTGGCTGCTGAAGAAGCAAAAGAAAAATATTTAATTGGCTT |
| R-AT10 | TAGCAAGCGCTCCTGCTGTGCCTAAGAATTCGCGGCGGCCCATGTGTACATTCCTCCTTCTCCTATAATGGTACC |
| F-aprE2-signalP- | GCCAAAGCTGTAATGGCTGAAAATTCTTAC |

| | |
|---|---|
| VF | |
| R-aprE2-signalP-VF | GGAAATACCTGTATCGAGGACAGCAAC |

**Table S5.** The signal peptide dataset compiled in this study.

| Dataset name | Filtered samples | Sec | Tat | No_SP | SP–MP pairs |
|---|---|---|---|---|---|
| UniRef50_sp | 3,570,658 | 3,496,344 | 74,314 | / | - |
| UniProtKB_SP | 208,051 | 193,530 | 14,521 | / | + |
| UniProtKB_UnSP | 201,036 | / | / | 201,036 | - |
| SP22 | 6,212 | 258 | 78 | 5,876 | - |
| SPSDB | 807 | 807 | / | / | - |
| SPE | 4,421 | 4421 | / | / | + |
| Signal6_set | 18,572 | 2582 | 365 | 15625 | - |

**Table S6.** Dataset Overlap Assessment.

| Model | Ref size | External set | Set size | Unit | overlap |
|---|---|---|---|---|---|
| Model2 | 409,087 | SP22 | 6,212 | N70 | 106/6,212 (1.71%) |
| Model2 | 409,087 | Signal6_Set | 18,572 | N70 | 995/18,572 (5.36%) |
| Model2 | 409,087 | External secretion panel | 586 | N70 | 0/586 (0%) |
| Model3 pretraining | 206,555 | SPE | 4,421 | SP–MP pair | 0/4,421 (0%) |
| Model3 fine-tuning | 210,976[1] | External secretion panel | 586 | SP–MP pair | 0/586 (0%) |

**Table S7.** Ablation analysis of the evolution-guided discrete diffusion model.

| Group | −1 site accuracy | −3 site accuracy | Top-5 | RR rate |
|---|---|---|---|---|
| Baseline | 0.6516 | 0.6693 | 0.9518 | 0.9005 |
| Baseline+A | 0.6786 | 0.6812 | 0.9522 | 0.9028 |
| Baseline+AB | 0.7327 | 0.7363 | 0.9744 | 0.9384 |
| Baseline+AC | 0.7043 | 0.6922 | 0.9518 | 0.8957 |
| Baseline+AD | 0.6745 | 0.6780 | 0.9523 | 0.9071 |
| Baseline+ABD | 0.7377 | 0.7379 | 0.9746 | 0.9463 |
| Baseline+ABC | 0.7648 | 0.7506 | 0.9745 | 0.9393 |
| Baseline+ABCD | 0.768 | 0.7520 | 0.9747 | 0.9530 |
| Baseline+ABCDE | 0.7531 | 0.7445 | 0.9746 | 0.9423 |

**Table S8.** Ablation analysis of the topology-aware multitask biological filter.

| Group | T1 Acc | T1 F1 | T2 Acc | T2 F1 | T3 Acc | T3 F1 |
|---|---|---|---|---|---|---|
| Base | 0.9934 | 0.9934 | N/A | N/A | 0.9297 | 0.9296 |
| Base+T2 | 0.9951 | 0.9951 | 0.9485 | 0.9369 | 0.9191 | 0.9187 |
| Base+Esm | 0.9973 | 0.9973 | N/A | N/A | 0.9622 | 0.9622 |
| Base+Esm+T2 | 0.9968 | 0.9968 | 0.954 | 0.9424 | 0.9584 | 0.9584 |

**Figures**

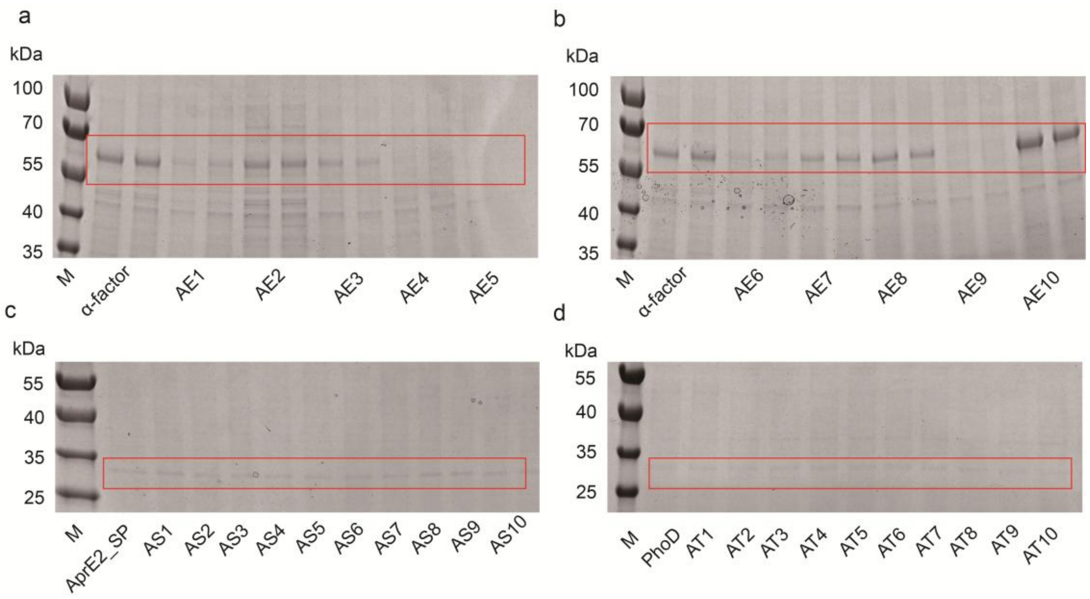


**Figure S1. SDS-PAGE gels illustrating signal peptide substitutions in eukaryotic and prokaryotic systems. (a, b)** Secretory expression of ApGA driven by α-factor and series AE signal peptides in *Pichia pastoris* X-33. **(c)** Secretion of AprE2 mediated by Sec-type signal peptides (AprE2_SP, AS1–AS10) in *Bacillus subtilis* WB600. **(d)** Secretion of AprE2 mediated by Tat-type signal peptides (PhoD, AT1–AT10) in *Bacillus subtilis* WB600.
